# Chromosome-scale pearl millet genomes reveal a *CARLACTONOIC ACID METHYL TRANSFERASE* as key determinant of strigolactone pattern and Striga susceptibility

**DOI:** 10.1101/2024.02.28.582441

**Authors:** Hendrik NJ Kuijer, Jian You Wang, Salim Bougouffa, Michael Abrouk, Muhammad Jamil, Roberto Incitti, Intikhab Alam, Aparna Balakrishna, Derry Alvarez, Cristina Votta, Guan-Ting Erica Chen, Claudio Martínez, Andrea Zuccolo, Lamis Berqdar, Salim Sioud, Valentina Fiorilli, Angel R de Lera, Luisa Lanfranco, Takashi Gojobori, Rod A Wing, Simon G Krattinger, Xin Gao, Salim Al-Babili

## Abstract

The yield of pearl millet, a resilient cereal crop crucial for African food security, is severely impacted by the root parasitic weed *Striga hermonthica,* which requires host-released strigolactones (SLs) for seed germination. Herein, we identified four SLs present in the Striga-susceptible line SOSAT-C88-P10 (P10), but absent in the resistant 29Aw (Aw). We generated chromosome-scale genome assemblies including four gapless chromosomes for each line. We found the Striga-resistant Aw lacks a 0.7 Mb genome segment containing two putative *CARLACTONOIC ACID METHYL TRANSFERASE1* (*CLAMT1*) genes. Upon transient expression, P10CLAMT1b produced methyl carlactonoate (MeCLA), an intermediate in SL biosynthesis. Feeding Aw with MeCLA resulted in the production of two P10-specific SLs. Screening a diverse pearl millet panel confirmed the pivotal role of the *CLAMT1* section for SL diversity and Striga susceptibility. Our results reveal a reason for Striga susceptibility in pearl millet and pave the way for generating resistant lines through marker-assisted breeding or direct genetic modification.

## Main text

Strigolactones (SLs) inhibit shoot branching and are released by plant roots into the rhizosphere to attract symbiotic mycorrhizal fungi (AM), particularly under phosphate (Pi) starvation, (Al-Babili and Bouwmeester, 2015; Fiorilli et al., 2019; Wang et al., 2023a). However, SLs also act as germination stimulants for root-parasitic weeds, such as *Orobanche* and *Striga spp.* (Yoneyama et al., 2010), posing severe agricultural problems worldwide (Parker, 2012). Moreover, the yield of pearl millet (*Pennisetum glaucum;* syn. *Cenchrus americanus*), the sixth most important grain globally (Satyavathi et al., 2021), primarily cultivated in subtropical regions, including Sub-Saharan Africa and India, is significantly affected by *Striga hermonthica* (Runo and Kuria 2018), which is considered one of the seven major threats to global food security (Pennisi, 2010).

After seed germination, Striga develops a haustorium that connects the emerging seedling to the host root. Host resistance to Striga comes in two forms. The first, post-attachment resistance, involves physical blocking, immune responses, or prevention of a vascular connection, and is triggered following haustorium attachment (Fishman and Shirasu, 2021; Dayou et al., 2021; Kavuluko et al., 2021). The second is a result of the amount and pattern of released SLs (pre-attachment) as has been proposed in studies regarding sorghum, rice and maize (Gobena et al., 2017; Ito et al., 2022; Li et al., 2023; Chen et al., 2023).

The wild pearl millet line 29Aw (Aw) from Niger exhibits resistance to Striga through both pre- and post-attachment mechanisms. In contrast, SOSAT-C88 P10 (P10) is a susceptible millet line derived from the SOSAT variety, which originates from a cross between the landraces Sauna and Sanio and has high yields in West Africa (Omanya et al., 2007; Dayou et al., 2021). To investigate the underlying resistance mechanisms, we first verified the contrasting phenotypes of the two lines under greenhouse conditions (Figure 1b, c, d). Root exudates from P10 induced higher Striga germination than those from Aw (Figure 1c), suggesting that P10 susceptibility may be related to released SLs. To test this assumption, we quantified the SLs exuded by both lines under Pi-deficient conditions using LC-MS/MS. Surprisingly, Aw exuded a significantly higher amount of the two canonical SLs orobanchol and orobanchyl acetate than P10 (Figure 1e). However, we identified four previously undescribed SLs of unknown structure in P10 exudates, based on the characteristic D-ring product ion peak at *m/z* 97.028. We named these SLs that were absent in Aw exudates as follows: Pennilactone1 (PL1) with a molecular ion formula C_20_H_23_O_6_ and a mass-to-charge ratio (*m/z*) at 359.14891 as a protonated positive ion ([M + H]^+^), Pennilactone2 (PL2) with a molecular ion formula C_20_H_25_O_7_ at *m/z* 377.15939 as [M + H]^+^, PL3 with a molecular ion formula C_20_H_23_O_7_; at *m/z* 375.14386 as [M + H]^+^, and PL4 with molecular ion formula C_19_H_32_O_12_ at *m/z* 452.19260 as [M + H]^+^, (Figure 1f, Supplementary figure 1, 2).

**Figure 1:**
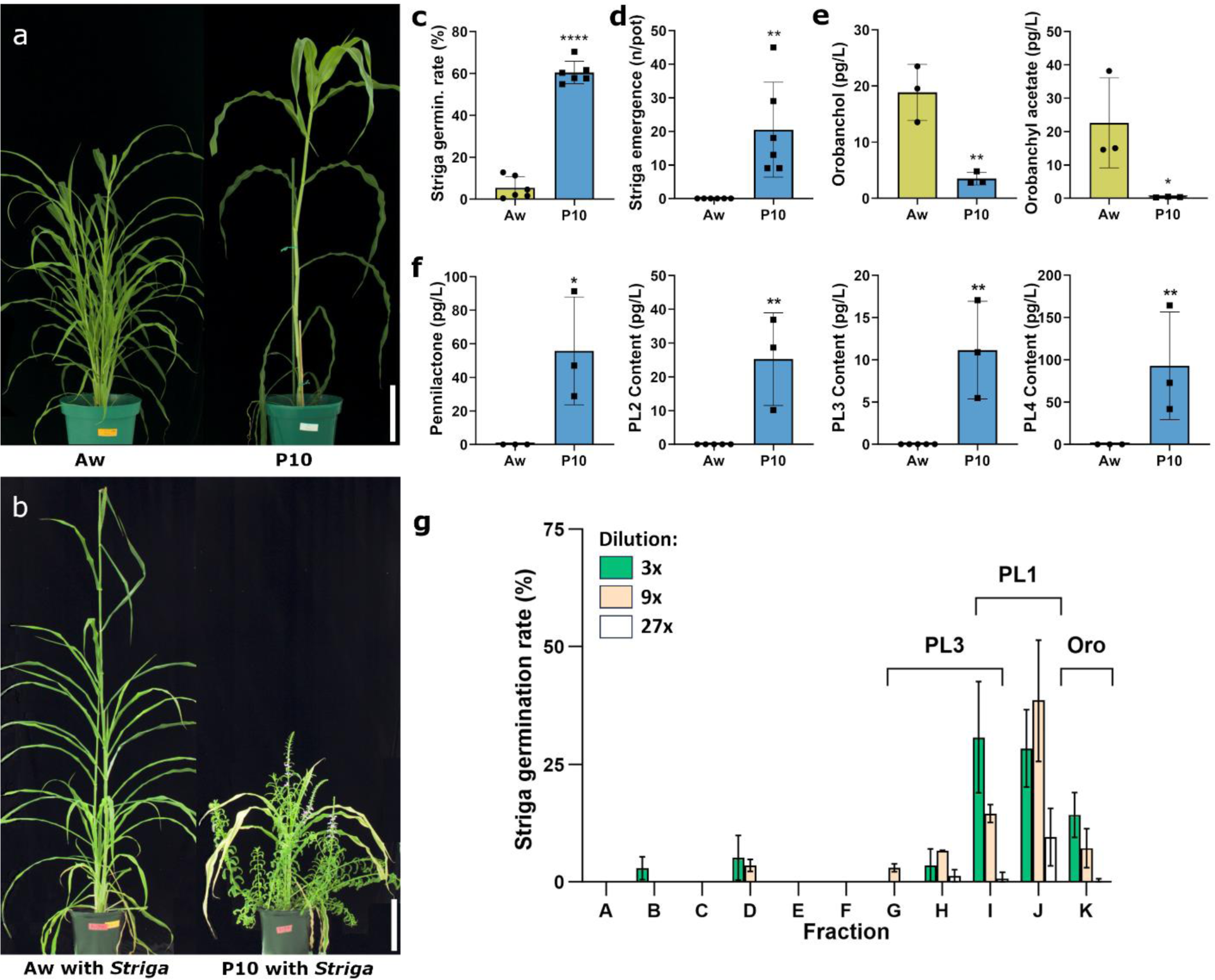
Aw and P10 are contrasting lines for Striga susceptibility and strigolactone production. **a**) The wild accession, Aw, was shorter and had several tillers, whereas the domesticated P10 was a tiller monoculm. **b**) Aw and P10 grown in Striga-infested soil. Aw exhibited no emerging Striga and was minimally affected in growth, while P10 was heavily infested by emerging Striga, severely impacting its growth and development. **c**) Less than 10% of the Striga seeds exposed to Aw’s root exudate germinated, whereas exposure to P10’s root exudate stimulated germination in over 70% of the Striga seeds. **d**) When grown in Striga-infested soil, P10 induced an average of 20 emerging Striga plants, while Aw induced none after 70 days. **e**) The root exudate of both Aw and P10 contained orobanchol and orobanchyl acetate, with significantly lower levels in P10. **f**) Four previously undescribed strigolactones were observed in P10’s root exudate but were never detected in Aw’s. **g**) Fractionation of SLs and subsequent assessment of individual fractions revealed that fractions G through J, containing PL and PL3, significantly induced Striga seed germination. Size bars indicate 20 cm. Error bars represent the mean ± s.d. Significant differences were tested using a two-tailed t-test (* P < 0.05, ** P < 0.01, *** P < 0.001, **** P < 0.0001).

Next, we evaluated the seed-germinating activity of the P10 SLs after fractionation using silica gel and detected significant activity in fractions enriched with PL1 and PL3 (Figure 1g). Confirming the SL identity of PL1, PL2, PL3 and PL4, we observed a significant decrease in their levels following zaxinone application, with a similar trend for its mimics, MiZax3 and MiZax5 that act as negative regulators of SL biosynthesis in rice (Supplementary figure 3a) (Wang et al., 2019; Wang et al., 2020). Additionally, the treatment with zaxinone growth regulators reduced Striga infestation, underscoring the role of SLs in P10 Striga susceptibility (Supplementary figure 3b,c). Recent studies have shown that canonical SLs with a tricyclic lactone (ABC-ring), such as orobanchol, do not have significant contributions to the inhibition of shoot branching/tillering, a function primarily attributed to non-canonical SLs (Wakabayashi et al., 2019; Ito et al., 2022; Chen et al., 2023; Cui et al., 2023; Wang et al, 2023b). Consistent with these findings, we observed contrasting tillering phenotypes in the two lines: Aw, lacking the four non-canonical SLs (PL1, PL2, PL3, and PL4), produced an average of 10 tillers, while P10 developed only one tiller under normal growth conditions (Figure 1a; Supplementary figure 4a). Our results suggest that the differing SL compositions of Aw and P10 likely account for their contrasting pre-attachment Striga resistance phenotypes.

To determine the genetic differences between Aw and P10 that underlie their contrasting SL compositions, we sequenced both genomes using PacBio HiFi Sequel II, allocating three SMRT cells to each, approximately 57-fold and 51-fold coverage, respectively. The contig N50 values obtained using hifiasm corresponded to 244 Mb for P10 and 284 Mb for Aw (Figure 2a). The assemblies were scaffolded using Omni-C technology. The final chromosome-scale genome assemblies both contained four gap-free chromosomes out of seven, with an order of magnitude fewer gaps than the previous best pearl millet assemblies, and total lengths of 1.915 Gb and 1.926 Gb for Aw and P10, respectively. We named and oriented the chromosomes of Aw and P10 based on the first reference genotype Tift 23D2B1-P1-P5 (Varshney et al. 2017). Guided by RNA sequencing and homology with genes from related grasses, gene structure annotation identified 38,920 high-confidence genes in Aw and 40,869 in P10, with BUSCO completeness scores of 92.9% for Aw and 93.3% for P10 (Supplementary table 1). The amounts of transposable element-related sequences were highly similar in the two accessions: 1.59 and 1.55 Gb corresponding to 82.90% and 80.50% of the total genome size for Aw and P10, respectively (Supplementary table 2). These values, which were larger and assigned in more detail than those reported by Varshney et al. in 2017 and Ramu et al. in 2023, aligned with the expectation that they represented at least 80% of the entire genome (Varshney et al., 2017). Our genome assemblies and findings are consistent with and improve upon previous pearl millet genome assemblies (Varshney et al., 2017; Yan et al., 2023; Ramu et al., 2023).

**Figure 2:**
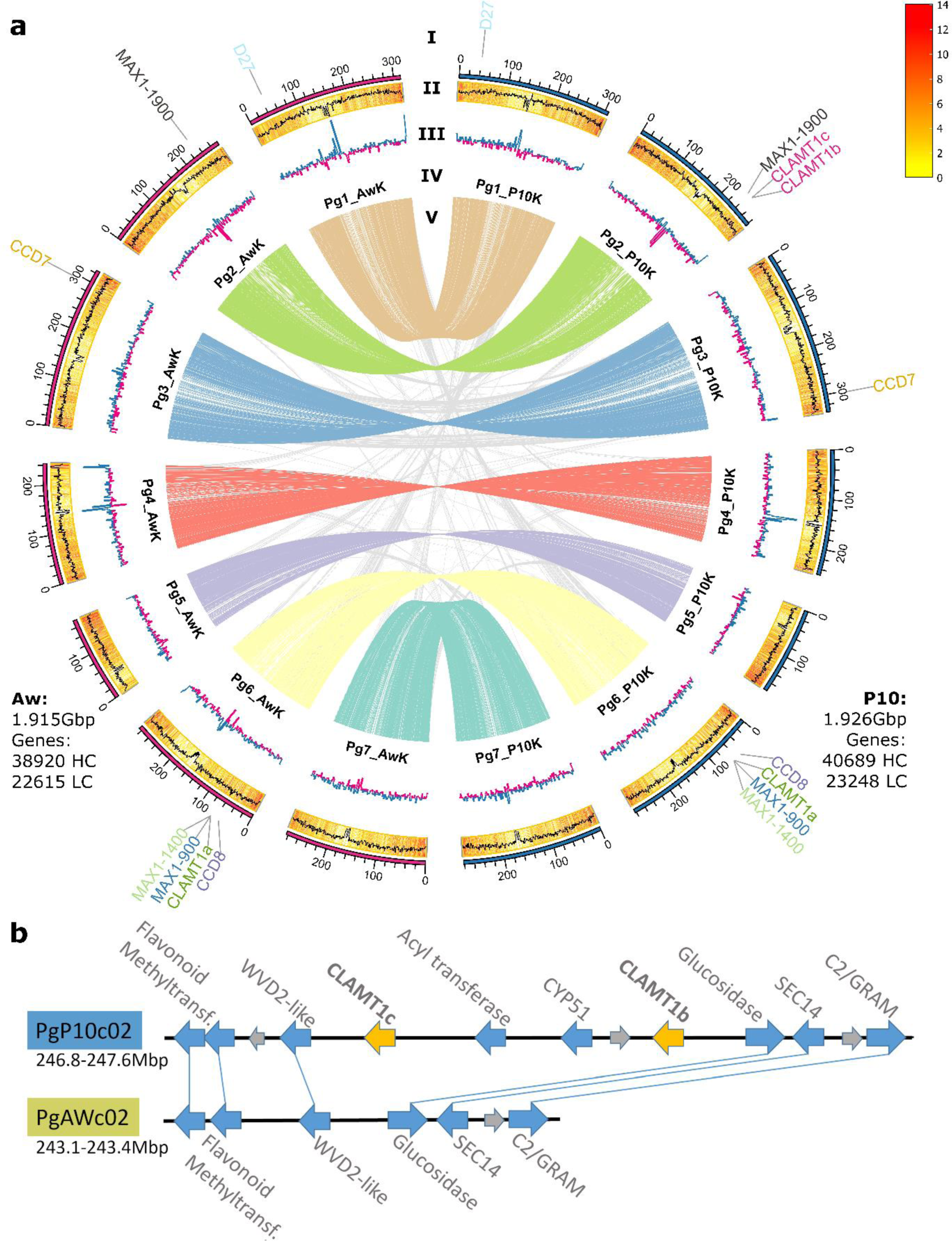
Genome assembly for Aw and P10. **a)** Circos plot of the genomes of Aw and P10. Layers from the edge to the center are as follows: I. Location of predicted strigolactone biosynthesis genes, II. GC content (line) and gene density (bars), III. GC skew, IV. Chromosome names, and V. Synteny plot. The insets provide the total assembly size and number of high-confidence (HC) and low-confidence (LC) genes. The synteny plot revealed no large-scale rearrangements between P10 and Aw. **b**) The 0.7 Mb section of chromosome 2, present in P10 but absent in Aw, extended from CLAMT1c to CLAMT1b and included CYP51 and an acyl transferase. The flanking regions exhibited strong synteny, as evidenced by three HC genes on each side of the region.

No large striking structural rearrangements existed between the genomes of Aw and P10 (Figure 2a). However, when homologs of SL biosynthesis genes were mapped to specific loci on Aw and P10 chromosomes (Figure 2a), we found that a smaller, 0.7 Mb segment of P10 chromosome 2 was absent in the Aw genome. This segment contained four predicted high-confidence genes, including two putative SL biosynthetic genes, *CARLACTONOIC ACID METHYLTRANSFERASE1b* (*CLAMT1b)* and *CLAMT1c* (Figure 2b, Supplementary figure 6).

Genome-wide association studies on Striga resistance in sorghum and maize, both closely related to millets, as well as in pearl millet itself, have surprisingly not yielded many strigolactone biosynthetic genes (Mallu et al., 2022; Kavuluko et al., 2021; Badu-Apraku et al., 2020; Rouamba et al., 2023). Although a comparison of the candidate genes from these studies with their homologs between Aw and P10 did not yield any results for pre-attachment resistance, it did indicate potential post-attachment resistance candidates (Supplementary table 5; Supplementary figure 7).

To identify candidate genes for SL biosynthesis in pearl millet, including the production of P10-specific SLs, we conducted a transcriptomic study on the root tissue of Aw and P10 under low Pi conditions, which stimulate SL biosynthesis, with and without treatment with the SL analog MP3, which was expected to modulate the transcript levels of SL biosynthetic genes. The total number of differentially expressed genes (DEGs) under both conditions was 5,833 in Aw and 6,890 in P10, with most being affected only by the low Pi condition (Supplementary figure 7a). The homologues of SL biosynthetic genes in pearl millet *PgD27* (*PgP10c0101G011332*), *PgCCD8* (*PgP10c0601G021829*), *PgMAX1*-*1400* (*PgP10c0601G022366*), *PgCYP706* (*PgP10c0401G051155*) and *PgCLAMT1b* (*PgP10c0201G045110*) (Supplementary figures 8, 9, 10) were all markedly upregulated in P10 under low Pi conditions (over 32-fold) and further by MP3 treatment. The canonical SL analog GR24 suppresses SL biosynthetic genes in rice, even under Pi deficiency (Haider et al., 2023). Therefore, the unexpected increase in expression observed here may be because MP3, a non-canonical SL analog, acts differently from GR24, or because SL biosynthesis is regulated differently in pearl millet compared to rice. We identified 30 and 42 genes induced by both treatments in P10 and Aw, respectively (Supplementary figure 7b; Supplementary table 3). Notably, *CLAMT1b* was absent in the Aw genome, as were *CLAMT1c* (*PgP10c0201G045096.1*), a *CYP51* homolog and a putative acyl transferase encoded by the 0.7 Mb fragment present in P10 (Figure 2b). CLAMT enzymes, involved in SL biosynthesis in rice and maize (Haider et al., 2023; Li et al., 2023), convert carlactonoic acid (CLA) into methyl carlactonoate (MeCLA) (Mashiguchi et al., 2022) in the formation of non-canonical SLs. In pearl millet, we identified three *CLAMT* genes with differing responses to phosphate starvation and MP3 treatment. While the *CLAMT1a (PgP10c0601G022359/PgAWc0601G022076.2)* transcript levels did not respond to either treatment, *CLAMT1c* was upregulated only under low Pi, and *CLAMT1b* transcripts increased strongly with both treatments (Supplementary figure 11), making it the primary candidate for the CLAMT function in pearl millet (Figure 3a).

**Figure 3:**
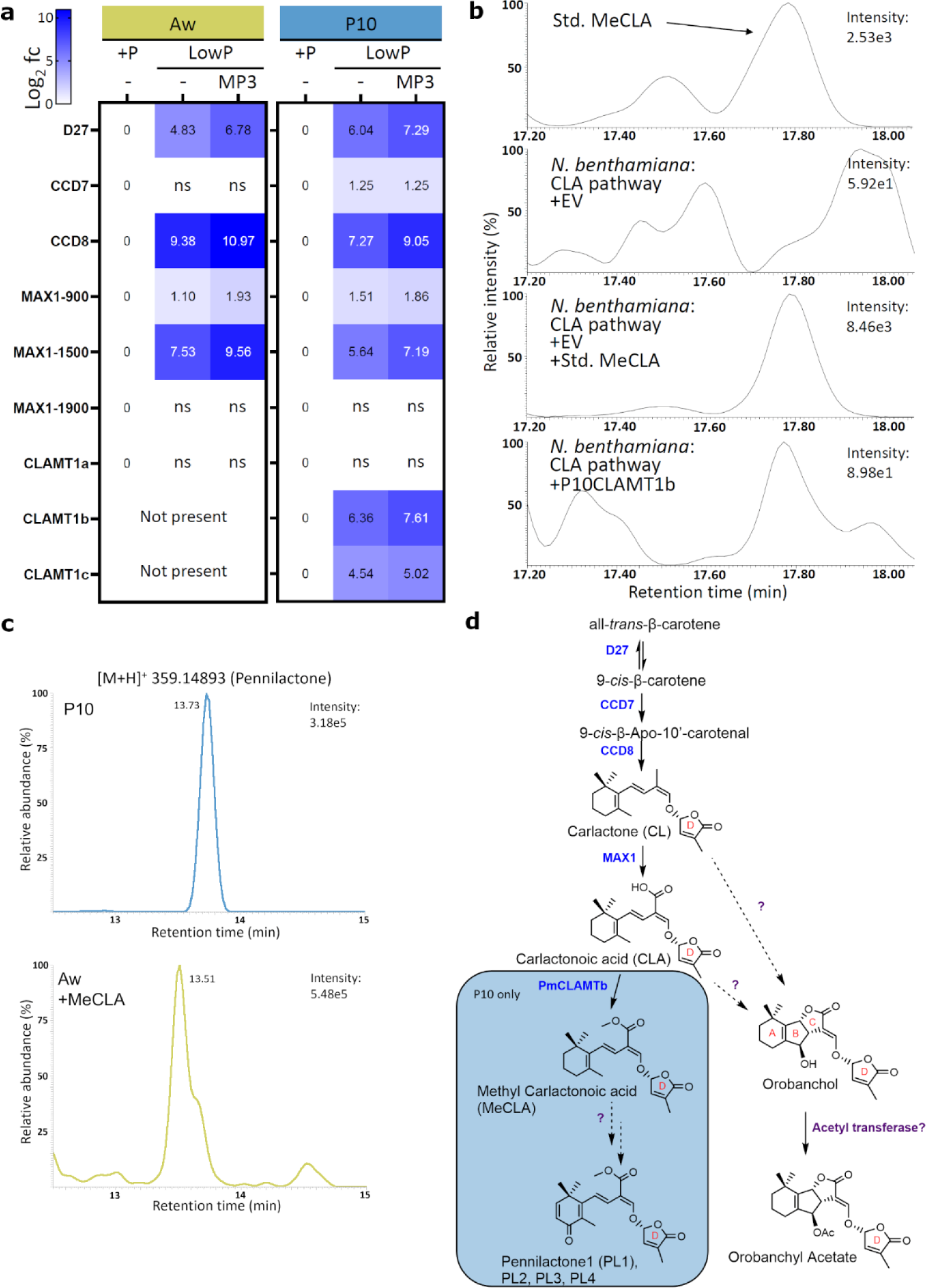
P10CLAMT1b produced methyl carlactonoate, which is a precursor to pennilactone. **a)** The expression of pearl millet homologs of known SL biosynthesis genes was generally upregulated in both Aw and P10 under low Pi conditions and even more so with the addition of the SL analog MP3. P10CLAMT1b was strongly co-expressed with PgD27, PgCCD8, and PgMAX1-1500, while P10CLAMT1c was induced more weakly. PgCLAMT1a was not significantly induced in Aw or P10. **b**) Recreating the SL biosynthetic pathway up to carlactonoic acid (CLA) through transient expression in *N. benthamiana* leaves produced no peak at the same retention time as the MeCLA standard. However, when P10CLAMTb was added to the experiment, a peak was observed, indicating that P10CLAMTb could convert CLA to MeCLA, as previously shown for OsCLAMT and AtCLAMT. **c**) Aw roots fed with MeCLA produced pennilactone (EIC: 359.14893; Retention time: 11.7 s), confirming that MeCLA is a precursor to pennilactone. **d**) The proposed SL biosynthesis pathway in pearl millet, where the boxed section is only present in P10 because of the presence of CLAMT1b. Consequently, only P10 produced downstream SLs, such as pennilactone1 (PL1), which can induce Striga seed germination when exuded from the roots, making P10 more susceptible to Striga infestation.

To test this assumption, we co-expressed PgCLAMT1b with OsD27, OsCCD7, OsCCD8 and AtMAX1, which give rise to CLA production, in tobacco leaves (Zhang et al., 2014). Subsequent LC-MS analysis revealed the formation of MeCLA, confirmed by comparison with a MeCLA standard based on the retention time and mass fragmentation patterns (Figure 3b; Supplementary figure 11). In contrast, co-expression of the CLA-forming enzymes with either CLAMT1a or CLAMT1c did not result in MeCLA formation (Supplementary figure 13), suggesting that CLAMT1b is crucial for MeCLA production in pearl millet. These results also suggest that MeCLA formation might be a necessary step for the biosynthesis of P10 non-canonical SLs, which could explain their absence in Aw. To test this hypothesis, we treated Aw seedlings grown under low Pi conditions with synthetic *rac*-MeCLA. Following feeding, we investigated the SL pattern of the root exudates and detected PL1 and PL2, as evidenced by the retention time, accurate mass, and fragmentation pattern (Figure 3c; Supplementary figure 14). Additionally, we proposed a putative structure for PL1 based on the mass difference and MS/MS fragmentations following MeCLA treatment (Supplementary figure 15). PL3 and PL4 were not detected, indicating that additional biosynthetic enzymatic activities, potentially absent in Aw, are required. This feeding experiment demonstrates CLAMT’s role in determining the SL pattern in pearl millet and explains the absence of pennilactone and PL2 in Aw. Based on chemical and genetic evidence, we established a model for the SL biosynthetic pathway in pearl millet. This model proposes that both Aw and P10 produce orobanchol and orobanchyl acetate; however, P10 has a unique pathway branch initiated by CLAMT1b that leads to MeCLA formation, which acts as the precursor of PL1 and PL2. The absence of this branch in Aw may account for the higher orobanchol and orobanchyl acetate content, as the metabolic flux is not divided, and there is no competition for the CL or CLA precursor.

To determine whether the association between *CLAMT1b* and Striga susceptibility was consistent across different pearl millet varieties, we examined a diverse set of previously sequenced pearl millet accessions (Yan et al., 2023) for the presence of this gene. We discovered that *CLAMTb* was consistently accompanied by *CLAMT1c*, *CYP51*, and an *Acyl transferase* in the same *CLAMTb*/c fragment (Supplementary table 4). We verified the presence of *CLAMTb/c* through genotyping (Supplementary figure 16) and identified it in two out of eight accessions. Subsequent SL analysis of the root exudates revealed the presence of PL1, PL3 and PL4, in both accessions containing *CLAMT1b/c* and PL2 in PI537069 only, unlike the exudates from the six accessions lacking the *CLAMT1b/c* region (Figure 4). This suggests that the biosynthesis of PL3 and PL4 might also rely on *CLAMT*. Consistent with the findings from P10, the root exudates of PI537069 and PI583800 exhibited significantly higher Striga seed-germinating activity than those from the six accessions lacking *CLAMT1b/c* region (Figure 4). These results demonstrate a clear link between *CLAMT1b/c*, the SL composition, and pre-attachment Striga susceptibility in this group of accessions. The ten pearl millet accessions did not share a common architecture or shoot phenotype (Supplementary figure 17a). We observed an inverse correlation between tiller number and Striga seed germination efficiency, with an *R^2^* of 0.70 for linear regression and a *P*-value of 0.0026. Although this correlation is significant, it is not highly predictive (Supplementary figure 17b,c). Tillering is a complex trait influenced by multiple factors, including SLs and others (McSteen, 2009), which can mask the impact of the absent *CLAMT1b* gene in several accessions.

**Figure 4:**
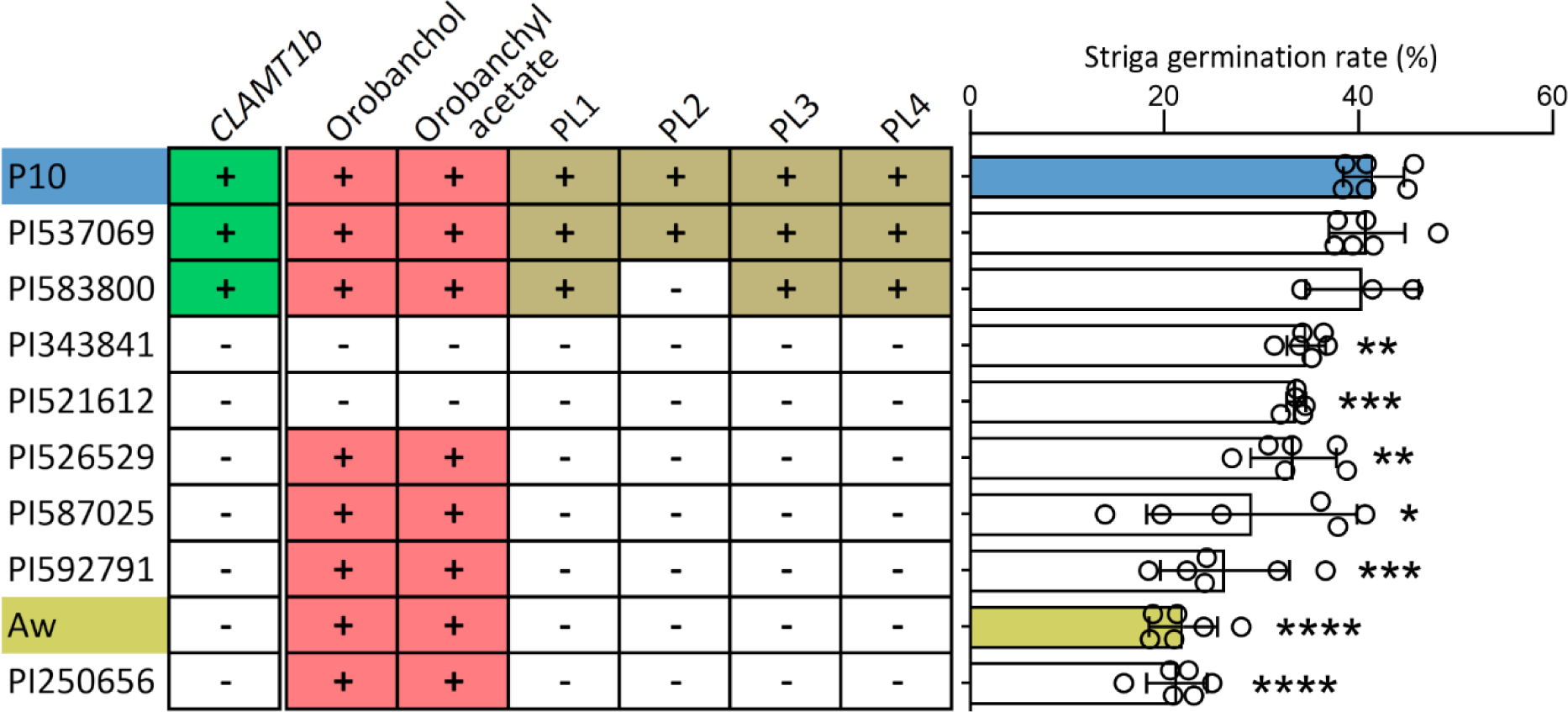
Pearl millet cultivars with the CLAMT1b gene exuded previously undescribed strigolactones and induced more Striga seed germination. The root exudates from P10, Aw, and eight other sequenced pearl millet accessions were analyzed for the presence of SLs and their ability to induce Striga seed germination. Only P10, PI537069, and PI583800 contained CLAMT1b. The lines are arranged in descending order of Striga germination rates. Error bars represent the mean ± s.d.; n = 6 biological replicates (except for PI583800, n = 3; and for PI521612, n = 5). Significant differences regarding P10 were assessed using a two-tailed t-test (* P < 0.05, ** P < 0.01, *** P < 0.001, **** P < 0.0001).

Analysis of resequencing data (Varshney et al., 2017) revealed that the *CLAMT* region was prevalent across a broad spectrum of cultivated pearl millet varieties and breeding stocks from various regions, as well as in certain wild varieties, indicating that our findings have global relevance (Supplementary figure 18). However, the existence of pearl millet accessions lacking the *CLAMT* fragment distributed across the same categories suggests that such accessions are viable and widely utilized. Consequently, the elimination of the *CLAMT* region to impart Striga resistance is unlikely to compromise the viability of an accession. Furthermore, mycorrhizal colonization, which is influenced by the composition of root exudate SLs, showed higher frequency and intensity values in Aw compared to P10 (Supplementary figure 19a) showing that removal of the *CLAMT* region does not reduce AM colonization and may even improve it. This finding was corroborated by molecular analyses of the expression of mycorrhiza-related marker genes (Supplementary figure 19b).

Altogether, we sequenced the genomes of agricultural pearl millet, SOSAT-C88 P10, susceptible to Striga, and 29Aw, a wild and resistant accession. The resulting platinum-grade, near gap-free genome assemblies, are the highest quality for pearl millet to date and can serve as the reference genome for global pearl millet research. Furthermore, we demonstrated that pre-attachment resistance to Striga in pearl millet depends on the types of SLs exuded by the roots. The critical gene for the presence or absence of the newly discovered SLs is *CLAMT1b*, located on a 0.7 Mb fragment of chromosome 2. This fragment is present in susceptible accessions, such as P10, and absent in accessions with higher resistance, such as Aw. This knowledge paves the way for the enhancement of agriculturally important pearl millet lines through marker-assisted breeding or direct genetic modification to confer Striga resistance.

## Materials and methods

### Genomic DNA extraction and sequencing

Aw and P10 seeds were germinated on half-strength Murashige and Skoog (MS) medium with agar and filter paper moistened with water respectively. The seedlings were grown in a soil/sand mixture under greenhouse conditions for 5 weeks, with the final 48 hours in darkness. From each plant, multiple leaf samples exceeding 1.5 g were harvested and immediately flash-frozen in liquid nitrogen. One leaf sample per plant was ground into a fine powder with mortar and pestle while still frozen, whereas a corresponding sample from the same plant was preserved at −80 °C.

High molecular weight DNA was extracted following the workflow for HMW DNA extraction for third-generation sequencing (Driguez et al., 2021) using the Genomics-Tip 100G kit (Qiagen), with the following modifications: For pearl millet, we used 1.5 to 2 g of ground tissue exceeding the standard recommendation of 1 g per four 100/G columns. Despite the protocol’s specific warnings against shaking the 50mL tubes during the lysis step, we found it necessary to occasionally disrupt tissue clots formation by shaking gently. The elution step extended to up to 4 hours instead of the 1 hour duration specified in the protocol.

Quality control of HMW DNA was performed using a FEMTO Pulse (Agilent) with the gDNA 165 kb kit (FP-1002-0275) and a separation time of 70 minutes. Both Aw and P10 samples contained a high amount of genomic DNA with a length above the cut-off of 50 kb. Quantification was conducted with a Qubit assay using the Qubit dsDNA BR assay kit (Thermo Fisher Scientific) yielding 288 ng/µl for Aw and 922 ng/µl for P10.

The HMW DNA was sheared to 20 kb and processed for PacBio HiFi sequencing by the KAUST Bioscience Core Lab. Samples were sequenced on a PacBio Sequel II using three SMRT cells each. The total throughput exceeded 30 Gb per SMRT cell for P10 and over 35 Gb per SMRT cell for Aw. The median read length was 15 kb per cell for P10 and 16 kb per cell for Aw.

### Omni-C library construction and sequencing

The Omni-C library was prepared using the Dovetail® Omni-C® Kit for plant tissues according to the manufacturer’s protocol. Chromatin was fixed with disuccinimidyl glutarate (DSG) and formaldehyde in the nucleus from dark-treated young leaves. The cross-linked chromatin was digested in situ with DNase I. After digestion, the cells were lysed with SDS to extract chromatin fragments, which were then bound to Chromatin Capture Beads. The chromatin ends were repaired and ligated to a biotinylated bridge adapter followed by proximity ligation of adapter-containing ends. After proximity ligation, the crosslinks were reversed, the associated proteins were degraded, and the DNA was purified. The purified DNA was then converted into a sequencing library using Illumina-compatible adapters. Biotin-containing fragments were isolated with streptavidin beads prior to PCR amplification. The two libraries were sequenced on an Illumina MiSeq platform to generate >214 and >269 millions 2 x 150 bp read pairs for Aw and P10, respectively.

### Genome assembly

PacBio HiFi reads were assembled using hifiasm (v17.6) (Cheng et al., 2021) with default parameters (https://github.com/chhylp123/hifiasm/) to generate primary contig assemblies. Subsequently, construction of the pseudomolecules was performed by integration of Omni-C read data using Juicer (v2; https://github.com/aidenlab/juicer)(Durand et al., 2016b) and the 3D-DNA pipeline (https://github.com/aidenlab/3d-dna) (Dudchenko et al., 2017). First, to generate the Hi-C contact maps for P10K and AwK genomes, Omni-C Illumina short reads were processed with juicer.sh (parameter: -s none --assembly). The resulting output file “merged_nodups.txt” and the primary assembly were then used to produce an assembly with 3D-DNA3 (using run-asm-pipeline.sh with the -r 0 parameter). Juicebox (v2.14.00) (Durand et al., 2016a) was employed to visualize the Hi-C contact matrix alongside the assembly and to manually curate the assembly. The orientation and order of each pseudomolecule were defined by dot-plot comparison using chromeister (https://github.com/estebanpw/chromeister) (Pérez-Wohlfeil et al., 2019) against the pearl millet genotype Tift 23D2B1-P1-P56. All the remaining contigs not anchored to the pseudomolecules were concatenated into “unanchored chromosomes”. The final Hi-C contact maps and assemblies were saved using run-asm-pipeline-post-review.sh from the 3D-DNA pipeline.

### RNA extraction and sequencing

Seeds of Aw and P10 descended from the sequenced individuals, were germinated as described above. The seedlings were transferred to 50 ml hydroponics tubes and grown in Hoagland solution modified in the following ways: no modifications, low phosphate (1% of normal P), low phosphate with MP3 (1.0 μM), and low phosphate with only acetone mock treatment. Treatment with MP3 and acetone mock were done only for the 6 hours before harvesting. The roots of the +P and lowP plants as well as 3 day old seedlings and a flowering inflorescence were sampled for Iso-Seq.

Roots and shoot stubs were collected separately from each hydroponic growth treatment for both Aw and P10, with samples pooled from three plants at each collection and four such biological replicates obtained.

All samples were flash-frozen in liquid nitrogen and ground to a fine powder in a sterilized mortar and pestle. Then 100 mg of the samples was separated for RNA extraction. RNA extraction was performed using an RSC 48 RNA extraction robot (Maxwell) and the Maxwell RSC Plant RNA kit (Promega). Although the RNA yield for some of the 64 samples was low, ranging from 36 ng/µl to 260 ng/µl as measured by NanoDrop, the purity was consistently high. The RNA integrity number (RIN) scores ranged from 8 to 10 for over 92% of the samples with an average RIN of 9.0.

The extracted RNA was submitted for to the KAUST Bioscience Core Lab for Iso-Seq sequencing. Samples from five different tissues for both Aw and P10 were tagged and multiplexed onto a SMRT cell each and sequencing on the PacBio Sequel II platform.

Samples for RNA-Seq were sent to Novogene (Singapore) for mRNA library preparation and 150 bp pair-end sequencing on Illumina’s NovaSeq 6000 platform, targeting a throughput of 12 Gb of data per sample. The returned data were consistently high quality, with the percentage of reads scoring a Phred value over 30 (indicating a base error below 0.1%) exceeded 90% for each sample.

### Transposable element identification and quantification

Transposable elements were identified searching the genome assemblies with the Extensive de-novo TE Annotator pipeline EDTA (version 2.0) (Ou et al., 2019) run under default settings. Due to the high incidence of false positivesin the prediction of helitrons, representatives of this class of TEs were removed from the final EDTA output. The TE library was then employed to mask the two genome assemblies and quantify the TE content using the tool RepeatMasker (http://www.repeatmasker.org/)) run under the default parameters (with the exception of the -qq option).

### Genome annotation

We annotated the two genomes using the MAKER pipeline v3.01.03 (Holt and Yandell, 2011). For a detailed breakdown of the genomes annotation, please refer to the supplementary materials (Supplementary figure 20) or the project’s GitHub page, https://github.com/mjfi2sb3/millet-genome-annotation. First, we prepared the necessary transcriptomic and homology data to inform and support the prediction in the MAKER workflow. We began by preprocessing the Iso-Seq data following PacBio’s recommended workflow using SMRT tools v11.0 (https://www.pacb.com/support/software-downloads/). The product of this step was a set of high-quality full-length isoforms for each submitted sample. Details regarding the preprocessing of RNA-seq data can be found in another section of this manuscript. For homology evidence, we incorporated the manually curated UniProt Swiss-Prot database (downloaded in Nov 2022) (Bairoch and Apweiler, 1997), along with published protein annotation for *Cenchrus americanus* (Varshney et al., 2017) and *Cenchrus purpureus* (elephant grass) (Yan et al., 2021). Additionally, we included NCBI annotations for *Setaria viridis* (green millet), *Setaria italica* (foxtail millet) and *Sorghum bicolor* (sorghum).

Subsequently, we processed the repeat-masked genome assemblies through the MAKER pipeline. The workflow calls an array of tools, including NCBI BLAST tools v2.2.28+ (Altschul et al., 1990), Exonerate v2.2.0 (Slater and Birney, 2005), Augustus v3.2.3 (Stanke and Waack, 2003) and tRNAscan-SE v2.0 (Lowe and Eddy, 1997). We ran MAKER’s workflow primarily with the default values except for alt_splice=1 and always_complete=1. Iso-Seq and RNA-seq transcripts were aligned to their respective assemblies using NCBI blastn, and these alignments were subsequently refined using Exonerate.

The protein evidence was aligned and refined using NCBI blastp and Exonerate, respectively. Subsequently gene structure prediction was performed using Augustus with the species parameter set to *Zea mays*. EST and protein hints were created using alignments obtained in the previous step. MAKER was then used to assess the predicted genes, correct some of the predictions, add isoform information and calculate quality scores (AED scores).

In the final step, we divided the predicted gene models into High Confidence (HC) and Low Confidence (LC) categories using four strategies: 1) based on EST evidence (MAKER’s quality index scores as well as alignment-based filtering); 2) annotating with the KEGG database (Kanehisa and Goto, 2000); 3) annotating using InterProScan v5 (Jones et al., 2014); and 4) annotating against the UniProt Swiss-Prot database.

### RNA-Seq data mapping onto P10 and Aw genomes

To map the RNA-Seq reads from each experiment using Spliced Transcripts Alignment to a Reference (STAR) software (Dobin et al., 2013), we first created an index of each of the P10 and Aw genome assemblies. For each RNA-Seq sample, the paired-end fastq data were then mapped on to the corresponding genome assembly using STAR with the option “–outSAMstrandField intronMotif” option. Subsequently, we assembled the transcripts for each RNA-Seq sample with StringTie (Pertea et al., 2015) using the BAM files generated in the previous alignment step. For each genome, we merged all transcripts from individual experiments using the StringTie merge option to produce a non-redundant set of transcripts.

### Determining differential expression

In all two-way comparisons we used the R Edger (Robinson et al., 2010) for differential expression analysis, with the default settings. We first filtered out genes having more than two replicates out of the total eight with a count per million (cpm) <=0.5. We performed the differential expression analysis using Fisher’s exact test and the p-values were adjusted for multiple testing using the Benjamini-Hochberg method.

### Gene identification and phylogeny

Homologs of known strigolactone biosynthetic pathway enzymes were identified through tblastn searches on the Aw and P10 genome assemblies using Persephone (persephonesoft.com). Phylogenetic trees of the protein families were constructed in Geneious v2023.2.1 (Biomatters) using muscle v5.1 based on the PPP alignment algorithm. The consensus tree was constructed using the neighbor-joining method, relying on the Jukes-Cantor model and was supported by 1000 bootstrap replicates.

### Screening diverse pearl millet accessions collection

We acquired a panel of 10 sequenced pearl millet accessions (Yan et al., 2023) from the U.S. National Plant Germplasm System. We screened the genomes of these accessions for the presence of the CLAMT region genes and the four flanking genes using tblastn, noting both presence and sequence similarity at the protein level (Supplementary table 4). Seedlings of the panel were genotyped through PCR using the Phire Plant Direct kit (Thermo Fisher Scientific) directly on leaf extracts. For two accessions, PI527388 and PI186338, we detected the presence of the CLAMT fragment through genotyping, despite its absence in their genome assemblies. Consequently, we excluded these two lines from subsequent experiments.

### Strigolactone collection, extraction, and measurements

Pearl millet seedlings were cultivated under controlled conditions with a day/night temperature of 28/22 °C. The seeds were surface-sterilized in a 50% sodium hypochlorite solution for 10 min and rinsed with sterile water. They were then placed in magenta boxes containing half-strength MS medium and allowed to germinate in darkness for 24 h. Following this period, they were incubated in a Percival chamber for 4 days. The germinated seedlings were then transferred into the soil for phenotyping, to sand for SL detection, or to a hydroponic system for *Striga* bioassays.

Analysis of SLs in root exudates was conducted using a previously published protocol (Wang et al., 2022). In summary, 1 L of root exudates, spiked with 0.672 ng of D6–5DS, was collected and applied to a C18-Fast Reversed-Phase SPE column (500 mg/3 mL; GracePure™) pre-conditioned with 3 mL of methanol and 3 mL of water. The column was then washed with 3 mL of water, and SLs were eluted with 5 mL of acetone. The SL fraction was concentrated to approximately 1 mL in an aqueous SL solution and subsequently extracted using 1 mL of ethyl acetate. Then, 750 µL of the SL-enriched organic phase was dried under a vacuum. The residue was reconstituted in 100 μL of acetonitrile:water (25:75, v/v) and filtered through a 0.22 μm filter for LC-MS/MS analysis.

For SL extraction from *N. benthamiana* leaf, samples were ground to a powder in liquid nitrogen using a mortar and pestle. About 300 mg of powder was weighed out and transferred to an 8 mL brown glass vial to which cold 2 mL ethyl acetate were added. After vortexing, sonication and centrifugation, the supernatant was transferred into an 8 mL glass vial. The pellet was extracted once more and the supernatants combined and dried in a SpeedVac. After drying, the residue was dissolved in 50 μL of ethyl acetate and 2 mL hexane. Further purification was performed using the Silica gel SPE column (500 mg/3 ml) preconditioned with 3 ml of ethyl acetate and 3 ml of hexane. After washing with 3 ml hexane, SLs were eluted in 3 ml ethyl acetate and evaporated to dryness under vacuum.

SL identification was performed using a UHPLC-Orbitrap ID-X Tribrid Mass Spectrometer (Thermo Fisher Scientific) equipped with a heated electrospray ionization source. Chromatographic separation was achieved using Hypersil GOLD C18 Selectivity HPLC Columns (150 × 4.6 mm; 3 μm; Thermo Fisher Scientific). The mobile phase comprised water (A) and acetonitrile (B), each containing 0.1% formic acid. A linear gradient was applied as follows (flow rate, 0.5 mL/min): 0–15 min, 25–100% B, followed by washing with 100% B, and a 3-min equilibration with 25% B. The injection volume was 10 μL, and the column temperature was consistently maintained at 35 °C. The MS conditions included: positive mode; spray voltage of 3500 V; sheath gas flow rate of 60 arbitrary units; auxiliary gas flow rate of 15 arbitrary units; sweep gas flow rate of 2 arbitrary units; ion transfer tube temperature of 350 °C; vaporizer temperature of 400 °C; S-lens RF level of 60; resolution of 120000 for MS; stepped HCD collision energies of 10, 20, 30, 40, and 50%; and a resolution of 30000 for MS/MS. The mass accuracy of identified compounds, with a mass tolerance of ± 5 ppm, is presented in Table 1. All data were acquired using Xcalibur software version 4.1 (Thermo Fisher Scientific).

SLs were quantified using LC-MS/MS with a UHPLC-Triple-Stage Quadrupole Mass Spectrometer (Thermo Fisher Scientific Altis^TM^). Chromatographic separation was achieved on a Hypersil GOLD C18 Selectivity HPLC Column (150 mm x 4.6 mm; 3 μm; Thermo Fisher Scientific), utilizing a mobile phase comprising water (A) and acetonitrile (B), each with 0.1% formic acid. The linear gradient was as follows (flow rate, 0.5 ml/min): 0–15 min, 25–100% B, followed by washing with 100% B, and a 3-min equilibration with 25% B. The injection volume was 10 μL, and the column temperature was consistently maintained at 35 °C. The MS parameters included: positive ion mode; H-ESI ion source; ion spray voltage of 5000 V; sheath gas flow rate of 40 arbitrary units; aux gas flow rate of 15 arbitrary units; sweep gas flow rate of 20 arbitrary units; ion transfer tube gas temperature of 350 °C; vaporizer temperature of 350 °C; collision energy of 17 eV; CID gas at 2 mTorr; and a Q1/Q3 mass with a full-width half maximum (FWHM) value of 0.4 Da. The characteristic Multiple Reaction Monitoring (MRM) transitions (precursor ion → product ion) were 347.14 → 97.02, 347.14 → 233.1, 347.14 → 205.1 for Oro; 389.15 → 97.02, 411.1 → 97.02, 389.15 → 233.1 for Oro Ace; 347.18→97.02, 347.18→287.1, 347.18→315.1, 347.18→329.14 for MeCLA; 299.09→158.06, 299.09→157.06, 299.09→97.02 for GR24; 359.14→ 97.02, 359.14→ 345.1, 359.14→ 299.1 for PL1; 377.15 →97.02, 377.15 →359.1, 377.15 →249.1 for PL2; 375.14 →97.02, 375.14 →343.1, 375.14 →247.1 for PL3; 452.19→97.02, 452.19→375.1, 452.19→315.1 for PL4.

### SL collection and fractioning

Analysis of SLs in root exudates followed the protocol by Wang et al. (2022). In summary, 1 L of collected root exudates was extracted using a C18-Fast Reversed-Phase SPE column (500 mg/3 mL; GracePure™), which had been pre-conditioned with 3 mL of methanol and 3 mL of water. The column was then washed with 3 mL of water, and SLs were eluted with 5 mL of acetone. The SL fraction was concentrated to approximately 1 mL of aqueous solution and then extracted with 1 mL of ethyl acetate. 750 μL of SL enriched organic phase was dried under vacuum. Concentrated SL extracts of root exudates obtained from 12 replicates (∼12L) were dissolved in 1.5 mL EtOAc/ 2 mL Hexane and subjected to silica gel column chromatography (SPE column 60g /50 mL) with a stepwise elution of Hexane/EtOAc (100:0–0:100, 10% step, 3 mL in each step) to yield 11fractions (A-K). 1 mL of each fraction was subjected to LC-MS analysis for monitoring the potential SLs and verify the Striga bioassay.

### Striga germination bioassays

The Striga germination bioassays were conducted following a previously described procedure (Jamil et al., 2023). In summary, Striga seeds were surface-sterilized with 50% diluted commercial bleach for 5 min. Then, they were dried and uniformly spread (approximately 50–100 seeds) on 9 mm filter paper discs made of glass fiber. Subsequently, 12 seed-laden discs were placed in a 9 cm Petri dish containing a Whatman filter paper moistened with 3.0 ml of sterilized Milli-Q water. The dishes were sealed with parafilm and incubated at 30 °C for 10 days for pre-conditioning. Post-conditioning, the Striga seeds were treated with SLs from root exudates of various pearl millet lines and incubated again at 30 °C for 24 h. Then, germinated and total seeds were scanned and counted using SeedQuant (Braguy et al., 2021), and the percentage of germination was calculated.

### Striga emergence under greenhouse pot conditions

The millet lines underwent Striga infection testing in pots within a greenhouse setting. Aproximately 2.0 L of blank soil, a mixture of sand and Stender soil, Basissubstrat, in a 1:3 ratio was placed at the base of an 8.0 L perforated plastic pot. Subsequently, approximately 40000 *Striga* seeds, equating to roughly 100 mg, were evenly distributed within a 5.0 L soil mixture and layered atop the blank soil in the pot. The *Striga* seeds within each pot underwent a pre-conditioning period of 10 days at 30°C with light irrigation maintained under greenhouse conditions. Following this, a single 10-day-old seedling was planted centrally in each pot. The millet plants were cultivated under standard growth conditions, with a temperature of 30°C and 65% RH. Striga emergence was monitored and recorded for each pot at 70 days post-millet sowing.

### Transient expression in *Nicotiana benthamiana* leaf

To generate pearl millet CLAMT plasmids for transient expression in *Nicotiana benthamiana*, the full-length cDNA of *CLAMT1b*, *CLAMT1a*, *CLAMT1c-*iso1, *CLAMT1c-*Iso2 (Supplementary table 6) were amplified by Phusion polymerase (New England Biolabs) from cDNA (*CLAMT1b*) or synthesized fragments (*CLAMT1a* and *CLAMT1c*; Azenta Life Sciences) using primers indicated in Supplementary table 7. The PCR products were purified and sequenced. Following Sanger sequencing, the gene sequences were amplified by using primers with suitable restriction enzyme sites. The resulting fragments were digested and ligated into the linearized entry vector pIV1A_2.1 which includes the CaMV35S promoter (www.pri.wur.nl/UK/products/ImpactVector/).

After sequence confirmation of the pIV1A_2.1 entry clones, Gateway LR clonase II enzyme mix (Invitrogen) reactions were performed to transfer the fragments into the pBinPlus binary vector (van Engelen et al.1995), generating p35S:PBIN-*CLAMT1b*, p35S:PBIN-*CLAMT1a*, p35S:PBIN-*CLAMT1c-*iso1 and p35S:PBIN-*CLAMT1c*-iso2.

Additionally, we cloned the Arabidopsis *Atmax1* and *Atclamt* cDNAs in the same binary vector; pBinPlus for transient expression in *N*. *benthamiana*.

The binary vector harboring various genes was introduced into *Agrobacterium tumefaciens* strain AGL0 via electroporation. Positive clones were cultured at 28°C at 220 rpm for 2 days in LB medium supplemented with 50mg/l Kanamycin and 35 mg/l Rifampicin. Cells were collected by centrifugation for 15 min at 4,000 rpm and room temperature. They were then resuspended in 10 mM MES-KOH buffer (pH 5.7) with 10 mM MgCl2 and 100 mM acetosyringone (49-hydroxy-3′,5′-dimethoxyacetophenone; Sigma) to achieve a final OD600 of 0.5. The suspension was incubated with gentle rolling at 22°C for 2–4 h. For various gene combinations, equal concentrations of Agrobacterium strains carrying different constructs were mixed, using strains with empty vectors to compensate for gene dosage in each combination. Additionally, an Agrobacterium strain containing a gene for the TBSV P19 protein was included to enhance protein production by inhibiting gene silencing. *N. benthamiana* plants were cultivated in soil pots in a greenhouse under a 14 h light/10 h dark cycle at 25 °C and 22 °C, respectively. Combinations of constructs in Agrobacterium were infiltrated into the leaves of 5-week-old *N*. *benthamiana* plants using a 1-ml syringe. Leaves at the same developmental stage were selected to reduce variability. For each gene combination, two to three leaves per plant were infiltrated, with three plants serving as individual biological replicates. The bacterial suspension was gently injected into the abaxial side of the leaf to ensure distribution throughout the entire leaf area. Six days post-infiltration, the leaves were collected for subsequent analysis.

### Analysis of resequencing data

We downloaded 1,036 sequence read archives (SRA) from NCBI SRA study SRP063925 and converted them to fastq files using sratools v3.0.7 (Sayers et al., 2022). We mapped the paired-end reads to our reference P10 assembly using the bwa-mem2 v2.2.1 mem subcommand with default parameters (Vasimuddin et al., 2019). We then extracted read mappings that fall within the region of interest (ROI). We estimated the mean coverage for the ROI and that of the harboring chromosome using the samtools v1.16.1 subcommand coverage with a minimum MAPQ score of 15 (Danacek et al., 2021). We calculated the coverage ratio (mean coverage of chromosome / mean coverage of ROI) as a proxy for the presence or absence of the ROI and plotted these ratios using Python. A detailed breakdown of the command workflow (Supplementary figure 19) is available on our GitHub page: https://github.com/mjfi2sb3/millet-genome-annotation.

### Mycorrhizal colonization of P10 and Aw

P10 and Aw were cultivated in sand and inoculated with approximately 1,000 sterile spores of *Rhizophagus irregularis* (DAOM 197198, Agronutrition, Labège, France). They received watering twice weekly, alternating between with tap water and a modified Long-Ashton (LA) solution containing 3.2 μM Na2HPO4·12 H2O.

All the plants were sampled at 45-days post-inoculation (dpi), corresponding to the late stage of mycorrhization. To evaluate the level of mycorrhization, we performed a morphological analysis according to Trouvelot et al. (1986). Moreover, we conducted qRT-PCR assays to assess the expression level of the fungal housekeeping gene (RiEF) and a fungal gene preferentially expressed in the intraradical structures (RiPEIP1) (Fiorilli et al., 2016), and two plant AM marker phosphate transporter genes (PtH1.9; EcPt4) (Ceasar et al., 2014; Pudake et al., 2017). We used alpha-tubulin (TUA; Saha et al., 2014) as the reference plant gene.

### Statistical Analysis

Data are represented as mean and their variations as SD. The statistical significance was determined by the two tailed unpaired Student t test or one-way ANOVA and Tukey’s multiple comparison test, using a probability level of P < 0.05. All statistical elaborations were performed using Prism 9 (GraphPad).

## Data availability

The Genome assemblies are available on the European Nucleotide Archive (ENA) under bioproject/study PRJEB71762. The same ENA study also contains: the genomic HiFi data, Omni-C data, RNA-Seq, Iso-Seq data, the assembled RNA-seq transcripts, and full-length Iso-Seq transcripts.

We have also hosted the above data on the DRYAD digital repository (https://doi.org/10.5061/dryad.nk98sf80k) along with the gene model predictions, repeat annotations and other useful information.

The Aw and P10 genomes were also uploaded to our platform at https://bioactives.kaust.edu.sa/Persephone and to the independently hosted platform at https://web.persephonesoft.com/. With data tracks for gene models, RNA-seq and Iso-Seq as well as BLAST search and synteny analysis functions.

The public resequencing data, SRA study SRP063925, were downloaded from NCBI: https://www.ncbi.nlm.nih.gov/Traces/study/?acc=SRP063925

## Acknowledgements

The authors would like to thank Prof. Steven Runo, Kenyatta University, Kenya for the P10 and Aw lines of great purity. Further thanks to Alexander Putra from the KAUST BioScience Core Lab for the primary sequencing, Vijayalakshmi Ponnakanti for greenhouse support, Luis River-Serna for technical support, and the other members of The BioActives Lab for miscellaneous support. We also thank the KAUST supercomputing laboratory (KSL) for providing resources and support; Dr. Helene Berges, Dr. Sandrine Arribat, Dr. William Marande and Dr. Stephane Cauet of INRAE, CNRGV French Plant Genomic Resource Center.

We thank Dr. Prakash Gangashetty and Dr. Rakesh Srivastava of ICRISAT-India for their support and for valuable discussions. We thank Prof. Harro Bouwmeester, Plant Hormones Biology, University of Amsterdam for valuable discussions and for providing plasmids for transient expression in *N. benthamiana* (pBIN-OsD27, pBIN-OsCCD7 and pBIN-OsCCD8), harboring the indicated cDNAs under the control of the CaMV 35S promoter.

This work was supported by baseline funding from King Abdullah University of Science and Technology and by the Bill & Melinda Gates Foundation (grant number OPP1136424) given to S. A.-B.

## Author information

These authors contributed equally: Hendrik NJ Kuijer, Jian You Wang, Salim Bougouffa

## Author affiliations

**The BioActives Lab, Biological and Environmental Sciences and Engineering (BESE), King Abdullah University of Science and Technology (KAUST), 23955-6900, Thuwal, Kingdom of Saudi Arabia** Hendrik NJ Kuijer, Jian You Wang, Muhammad Jamil, Aparna Balakrishna, Derry Alvarez, Guan-Ting Erica Chen, Lamis Berqdar, Salim al-Babili

**Center for Desert Agriculture, King Abdullah University of Science and Technology (KAUST), Thuwal, 23955-6900, Saudi Arabia**

Hendrik NJ Kuijer, Jian You Wang, Michael Abrouk, Muhammad Jamil, Aparna Balakrishna, Derry Alvarez, Guan-Ting Erica Chen, Lamis Berqdar, Andrea Zuccolo, Rod A Wing, Simon G. Krattinger, Salim al-Babili

**Plant Science Program, King Abdullah University of Science and Technology (KAUST), Thuwal, 23955-6900, Saudi Arabia**

Michael Abrouk, Andrea Zuccolo, Rod A Wing, Simon G. Krattinger

**Computational Bioscience Research Center, Computer, Electrical and Mathematical Sciences and Engineering Division, King Abdullah University of Science and Technology (KAUST), 23955-6900, Thuwal, Kingdom of Saudi Arabia**

Salim Bougouffa, Roberto Incitti, Intikhab Alam, Xin Gao

**Department of Life Sciences and Systems Biology, University of Torino; Viale Mattioli 25, Torino, 10125, Italy**

Cristina Votta, Valentina Fiorilli, Luisa Lanfranco

**Universidade de Vigo, Facultade de Química and CINBIO, 36310 Vigo, Spain**

Claudio Matrtinez, Angel R. De Lera

**Crop Science Research Center, Sant’Anna School of Advanced Studies, Pisa, Italy**

Andrea Zuccolo

**Analytical Chemistry Core Lab, King Abdullah University of Science and Technology (KAUST), Thuwal, 23955-6900 Saudi Arabia**

Salim Sioud

## Author contributions

The study was conceived by S.A-B.

Research designed and concept proposed by J.Y.W. and S.A-B.

Genomic DNA extraction by H.N.J.K., genome assemblies by M.A., genome annotations by S.B., repeat analysis by A.Z.

Conceptualization of genome sequencing: M.A., S.G.K., R.A.W.

RNA-Seq by H.N.J.K. with L.B., mapping by I.A., data processing to DEG tables by R.I.

CLAMT fragment analysis in resequencing data by S.B.

Interpretational bioinformatics by H.N.J.K.

Discovery of new SLs and fractioning by J.Y.W. MeCLA synthesis by C.M and A.R.d.L.

Chemical analysis, treatments and feeding by J.Y.W with G.T.C. and S.S.

Striga bioassay and pearl millet phenotyping by M.J., J.Y.W., L.B. and H.N.J.K.

Transient expression by A.B. and D.A. with H.N.J.K.

Mycorrhiza colonization analysis by C.V., V.F. and L.L.

Manuscript written by H.N.J.K., J.Y.W. and S.A-B. with input from all authors.

## Corresponding author

Correspondence to Salim al-Babili

