## Supplementary materials for "Chromosome-scale pearl millet genomes reveal a *CARLACTONOIC ACID METHYL TRANSFERASE* as key determinant of strigolactone pattern and Striga susceptibility"

**This file includes:**

**Supplementary Figure 1-21**

**Supplementary Table 1-7**

**References**

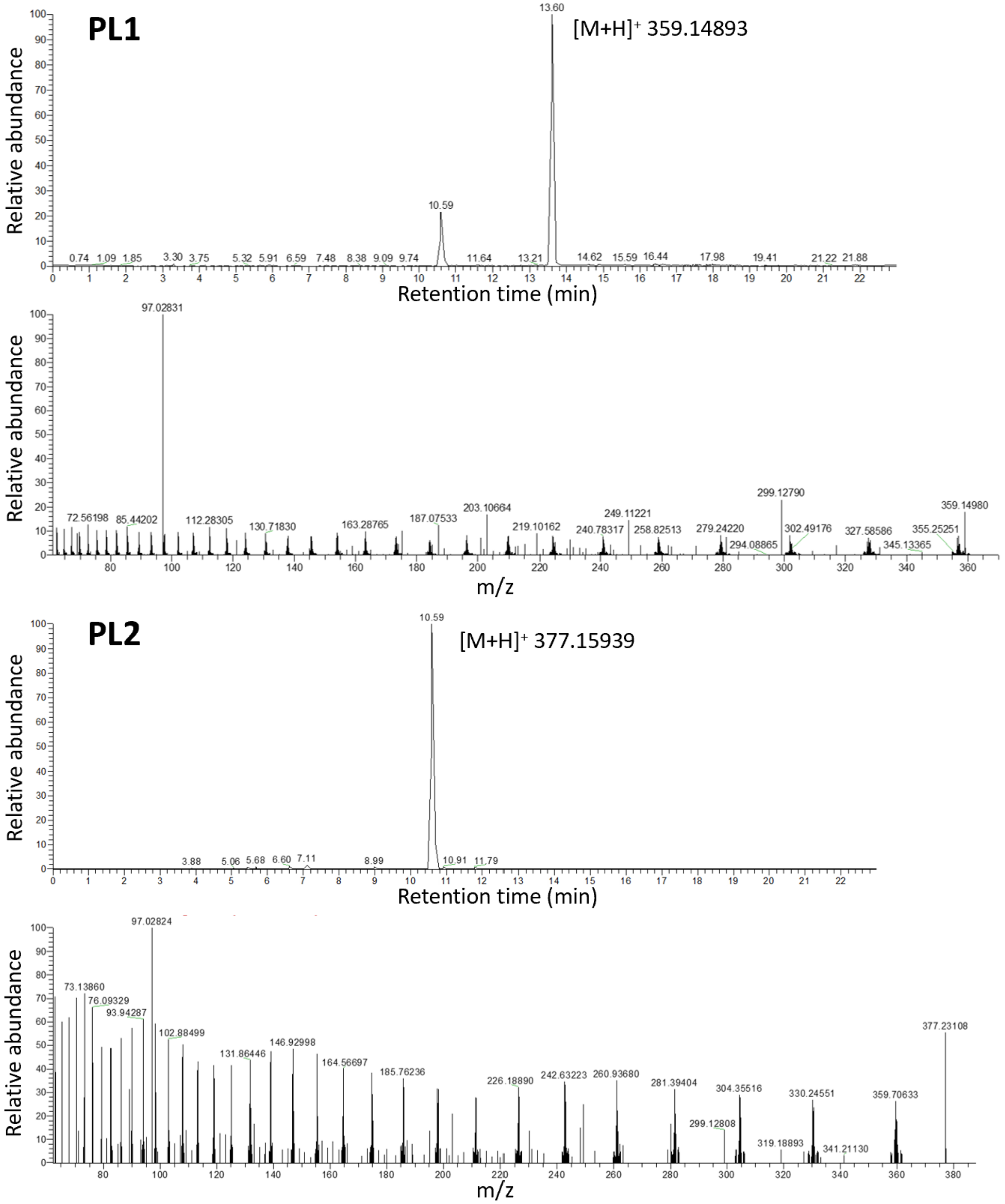

**Supplementary Figure 1. The chromatography of pennilactone (PL) and PL2 identification from P10 root exudate.**

**a** EIC chromatograms of PL1 (m/z 359.14893 $[M+H]^+$ in positive mode; retention time 13.60) and MS/MS fragmentation of PL with the characterized D-ring at 97.02831. **b** EIC chromatograms of PL2 (m/z 377.15939 $[M+H]^+$ in positive mode; retention time 10.59) and MS/MS fragmentation of PL2 with the characterized D-ring at 97.02824.

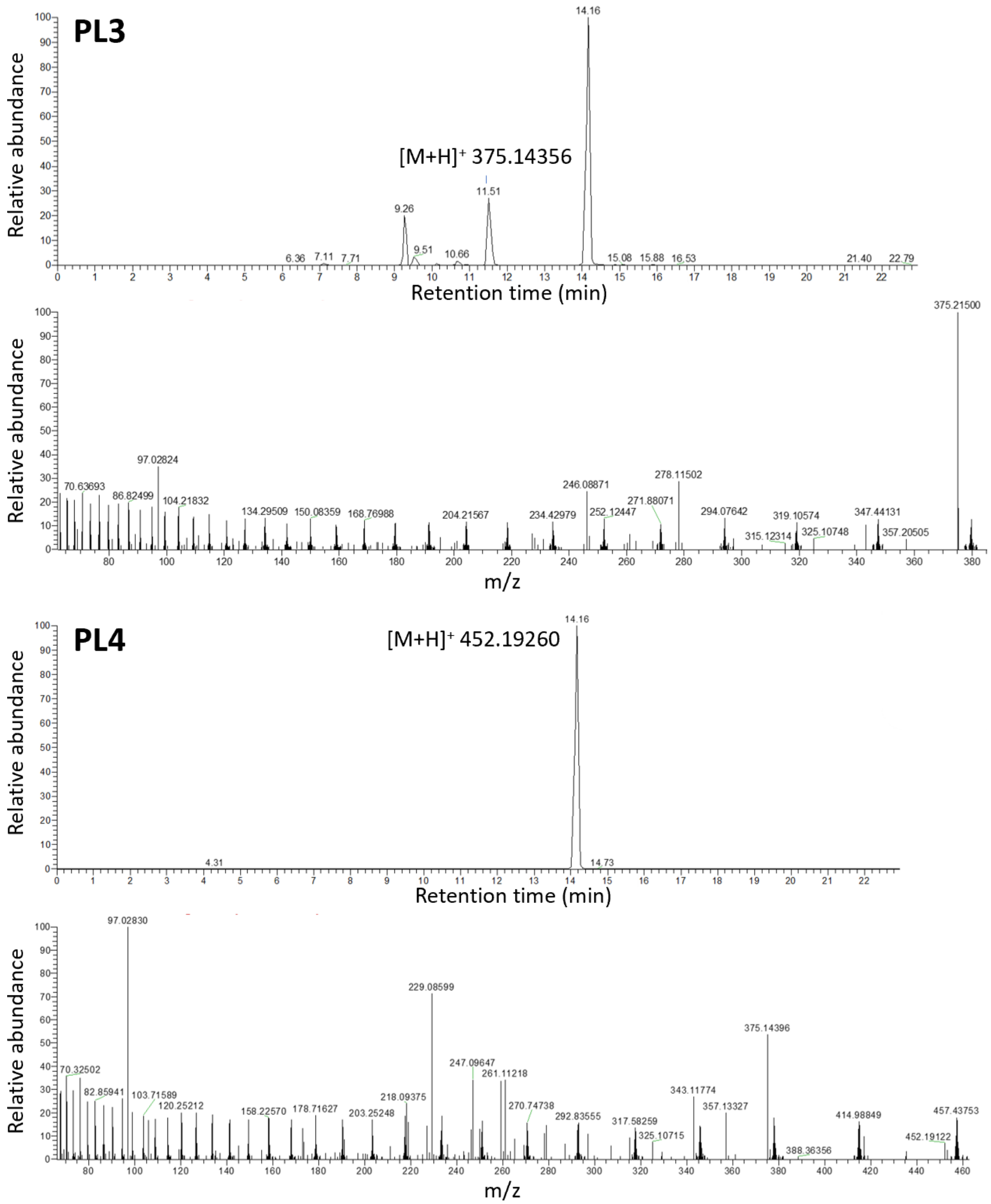

**Supplementary Figure 2. The chromatography of PL3 and PL4 identification from P10 root exudate.**

**a** EIC chromatograms of PL3 (m/z 375.14356 $[M+H]^+$ in positive mode; retention time 11.51) and MS/MS fragmentation of PL3 with the characterized D-ring at 97.02824. **b** EIC chromatograms of PL4 (m/z 452.19260 $[M+H]^+$ in positive mode; retention time 14.16) and MS/MS fragmentation of PL4 with the characterized D-ring at 97.02830.

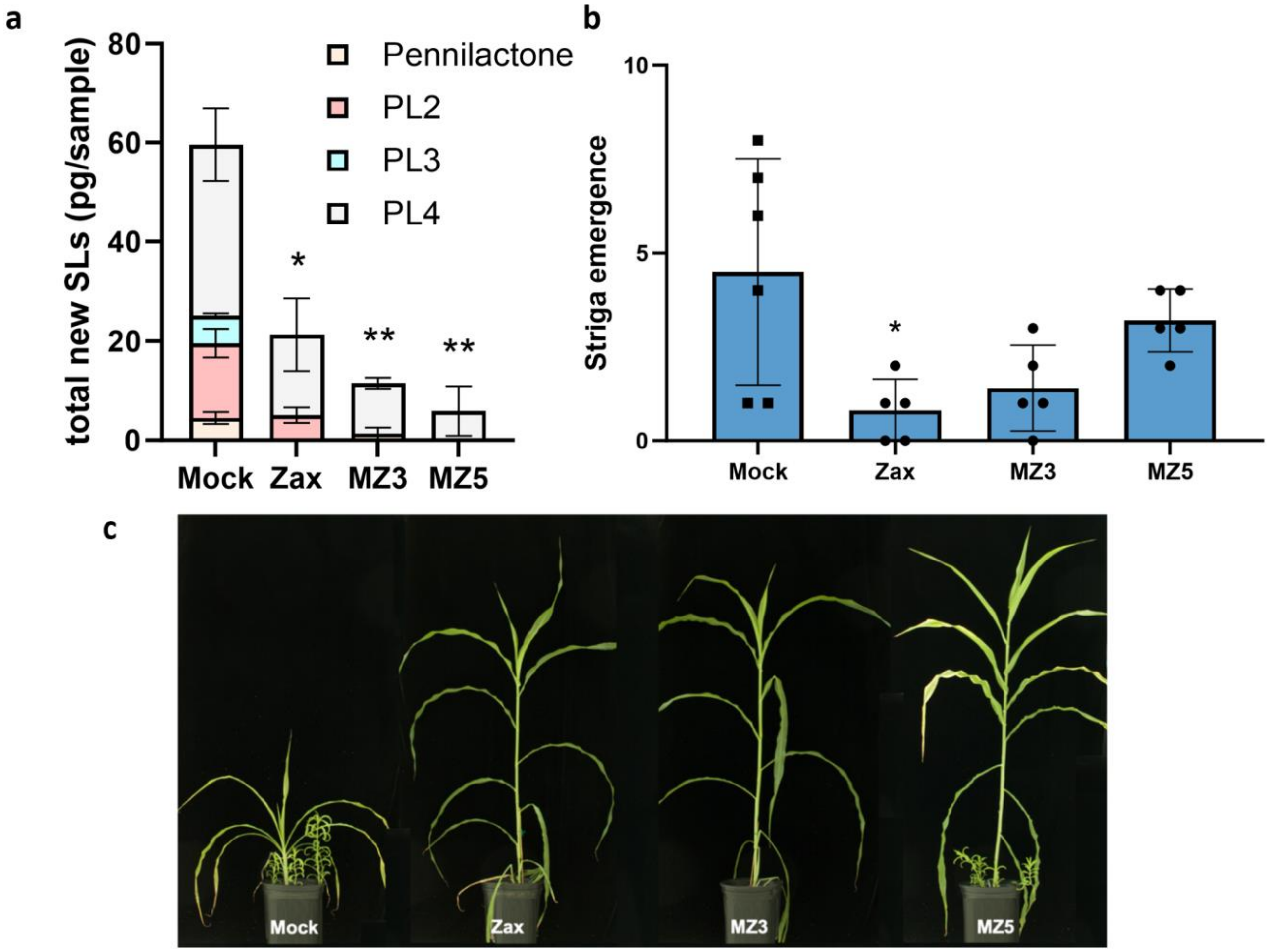

**Supplementary Figure 3. Zaxinone and MiZax suppress the production of new SLs in P10, reducing susceptibility to Striga.**

**a** Treatment of P10 with zaxinone (Zax) or MiZax3 (MZ3) or MiZax5 (MZ5) significantly reduces the amount of the newly discovered SLs exuded from the roots, showing that pennilactone, PL2, PL3 and PL4 act like SLs in respect to zaxinone. **b** Striga emergence is significantly reduced when P10 is treated with 0.25 µM of zaxinone, while treatment with MiZax3 and MiZax5 follows the same trend, but does not reach significance in this experiment. **c** Zaxinone and MiZax treatments partially rescues the Striga induced phenotype of P10, so zaxinone reduces the susceptibility of P10 to Striga; likely by suppressing the biosynthesis of the four new SLs. Significant differences to mock treatment were tested using ROUT (Q=5%) to discard outliers followed by a two-tailed t-test (* $P < 0.05$, ** $P < 0.01$, *** $P < 0.001$, **** $P < 0.0001$).

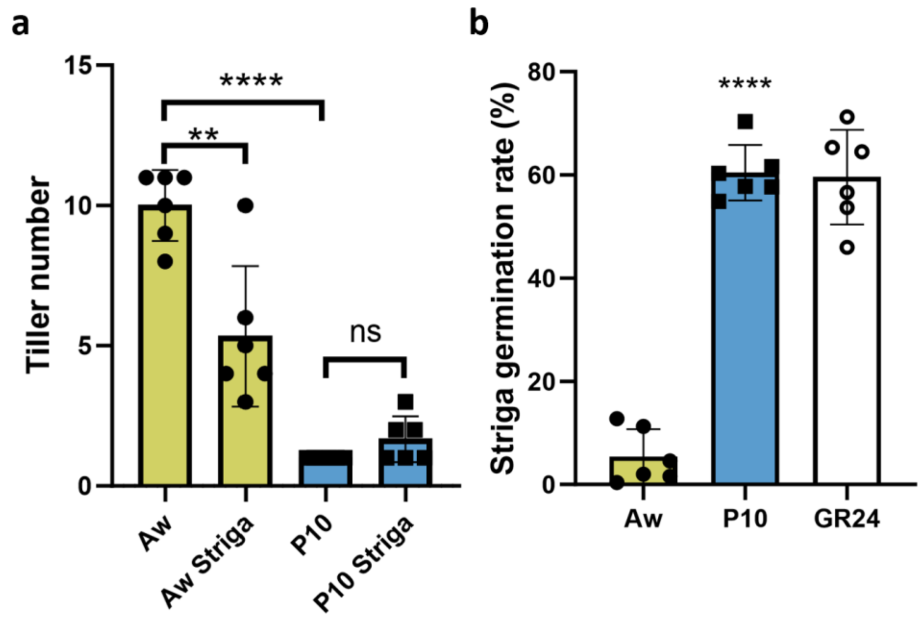

**Supplementary Figure 4. Additional phenotyping of Aw and P10 under greenhouse conditions.**

**a** The tiller number of Aw is reduced from an average of 10 under normal growth conditions to 5 when growing in Striga infested soil. P10 does not make any tillers in normal conditions, while it sometimes makes additional tillers when grown in striga infested soil, but this increase is not significant. **b** The Striga seed germination rate induced by P10 root exudate is not only significantly higher than that of Aw as also shown in main figure 1, but it reaches the rate of the positive control GR24, an artificial SL analog. This indicates the maximum germination rate of the seed batch under the circumstances of this experiment is reached and the contrast between Aw and P10 may be even greater than depicted here.

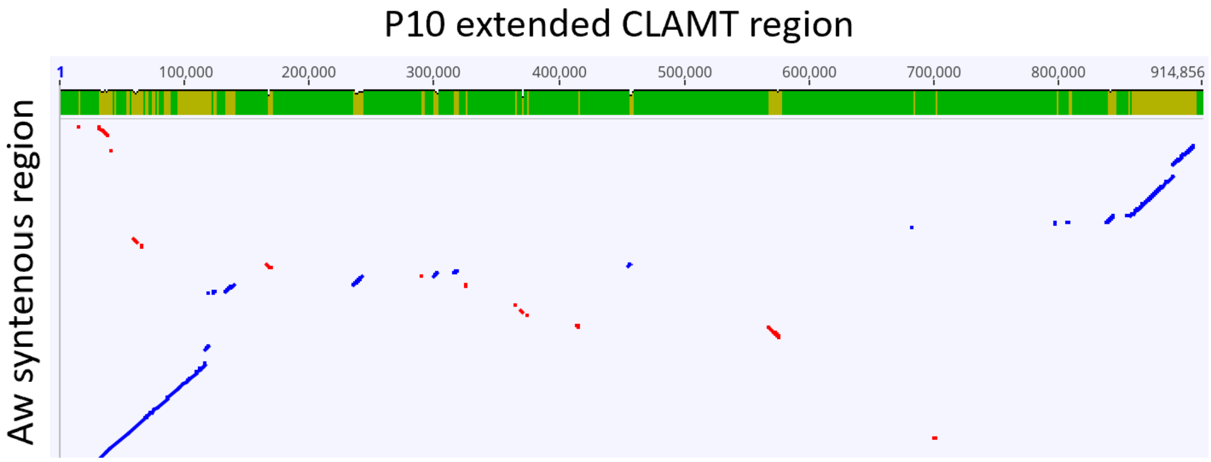

**Supplementary Figure 5**. **Alignment of the *CLAMT* region and the flanking sequences on chromosome 2 of P10 and Aw**.

The *CLAMT* region on P10 is 0.7Mbp and contains four HC genes, while the equivalent region in Aw is only 0.1Mbp and has no predicted genes. Figure created with LASTZ 7.0.3 plugin for Geneious Prime.

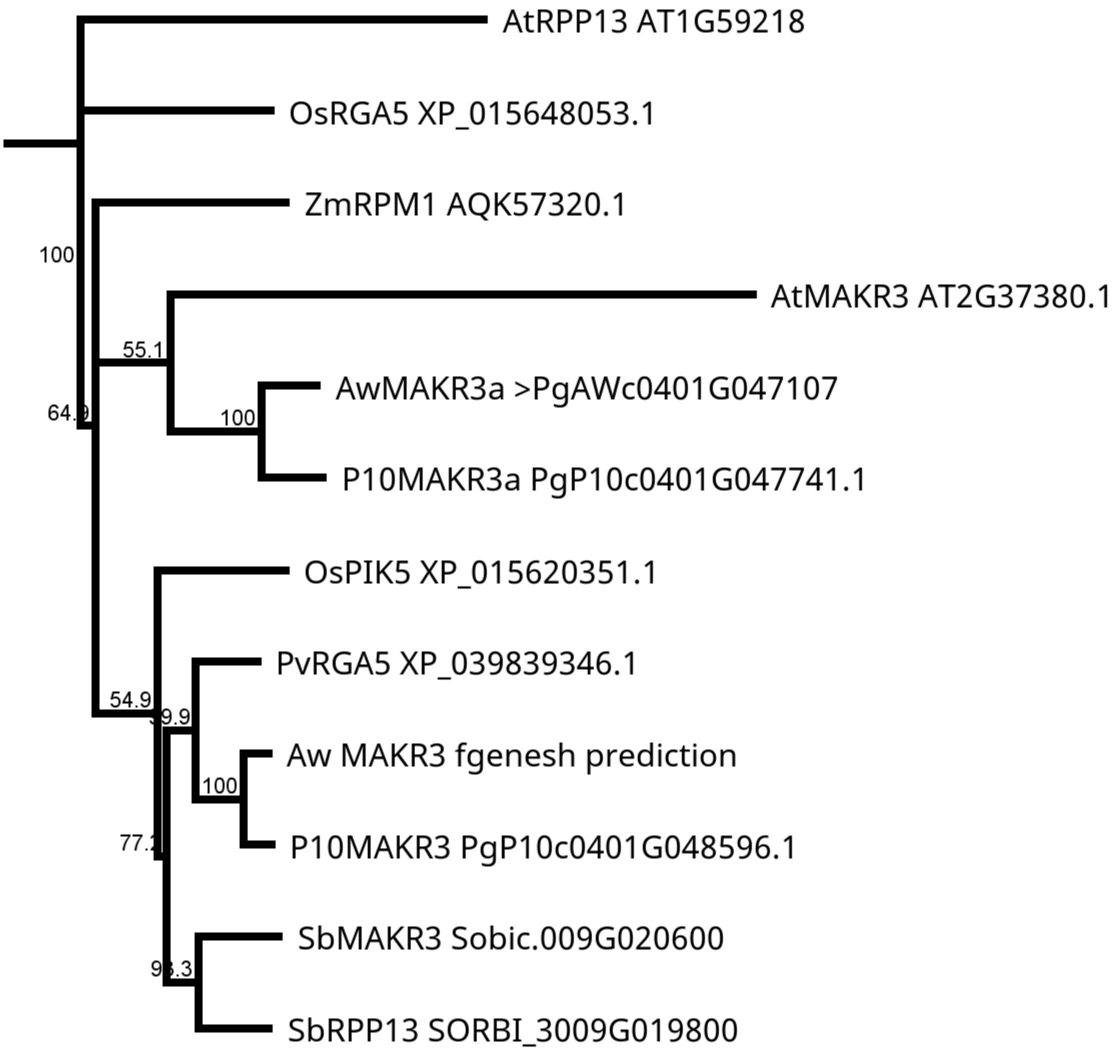

**Supplementary Figure 6**. **Phylogenetic tree for the MAKR3 gene family**.

*SbMAKR3* is a candidate gene for Striga resistance in sorghum found through GWAS (Mallu et al., 2022). We named the closest homologs in pearl millet *P10MAKR3* and *AwMAKR3*, but found they are CC-NB-LRR family genes and the link to *AtMAKR3* was erroneous. Both pearl millet MAKR3s and the sorghum MAKR3 contain the CC, NB and LRR domains, while the passing resemblance to *AtMAKR3* is spread out over the genes in small blocks. The automated annotation of *AwMAKR3* split the gene over two gene numbers, but a targeted manual annotation with Fgenesh (Softberry) shows the gene is present and intact in Aw.

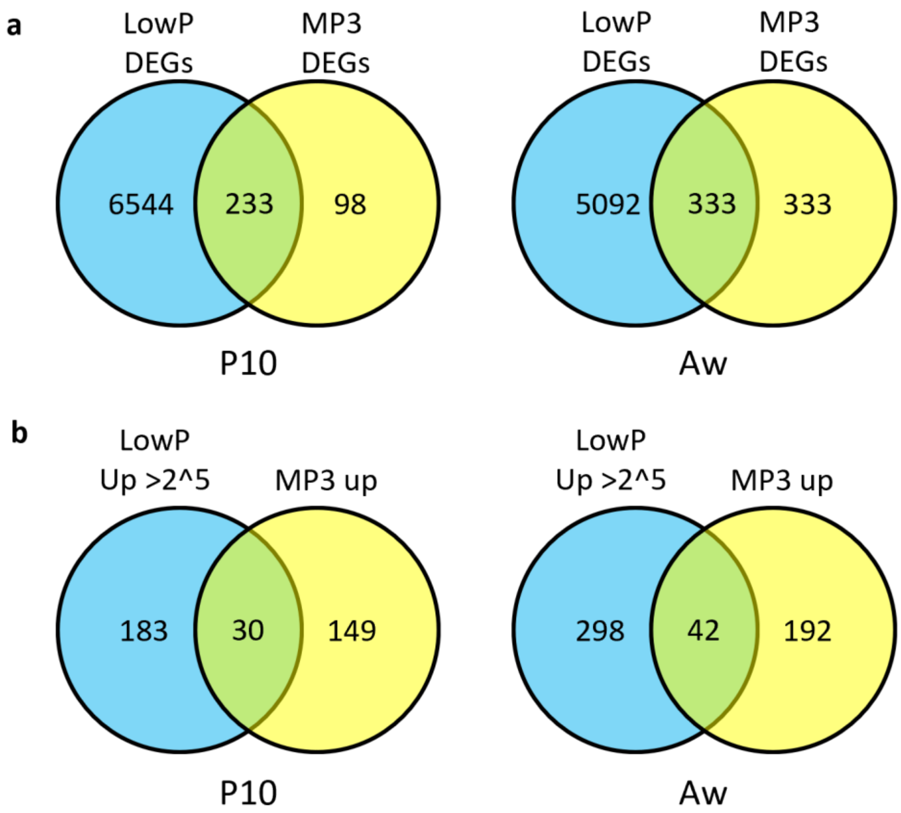

**Supplementary Figure 7. Differentially expressed genes under low phosphate and MP3 treatment.**

**a** Differentially expressed genes in both P10 and Aw roots when exposed to low phosphate are plentiful, while MP3 (an artificial SL analog) affects much fewer genes. The number of genes affected by both is 233 for P10 and 333 for Aw. **b** Narrowing down the selection to genes strongly upregulated by phosphate starvation (over 2^5 or 32 fold) and upregulated by MP3 leaves only 30 genes for P10 and 42 for Aw. The resulting list for P10 still includes predicted SL biosynthesis genes *D27*, *CCD8*, *MAX1-1400* and *CLAMT1b* (Supplemental table 3).

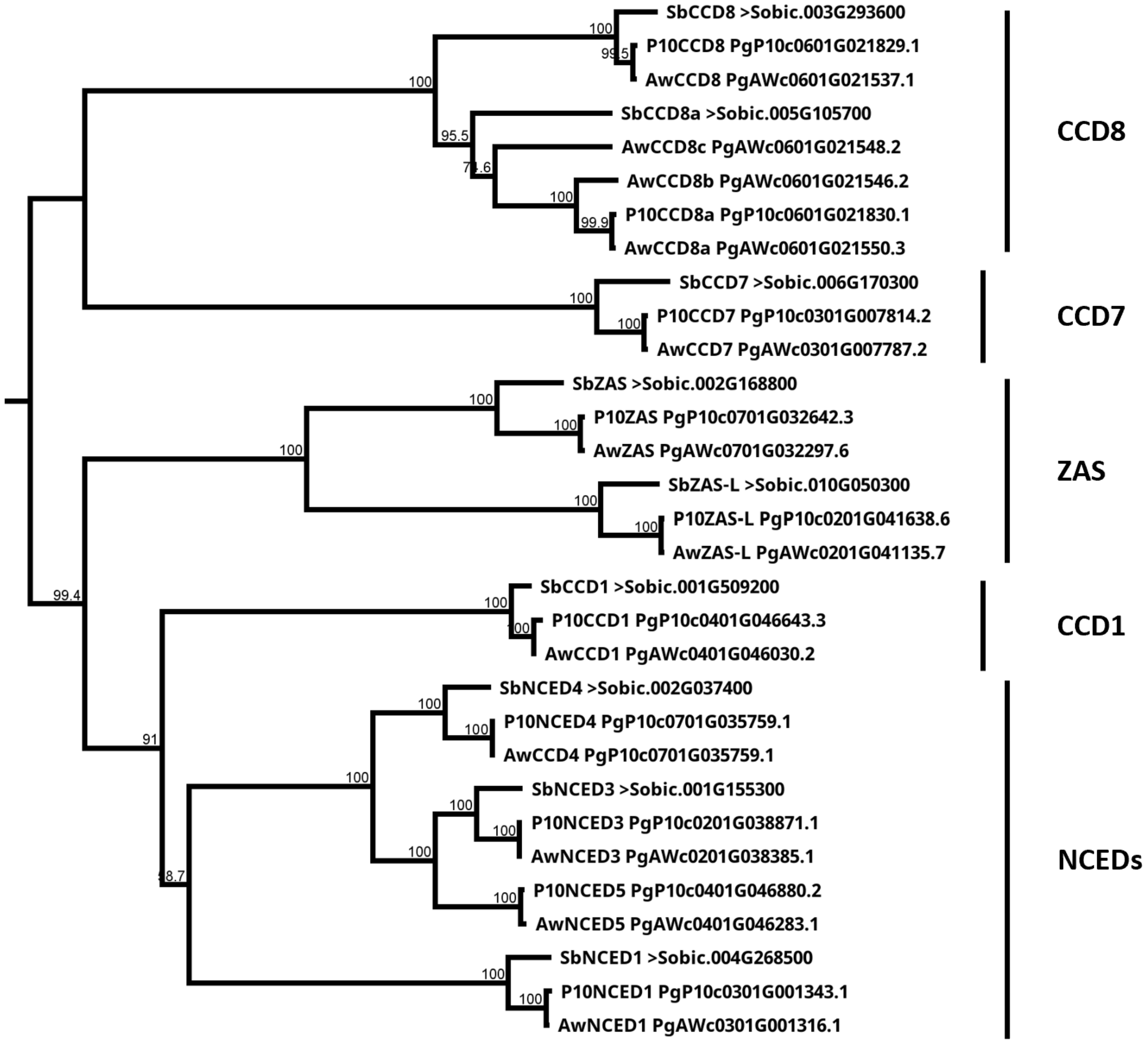

**Supplementary Figure 8**. **Pearl millet CCDs phylogenetic tree**.

Most members of the PgCCD family have a one-on-one homologous counterpart in sorghum. The only exception is CCD8, where P10 has one additional CCD8-like gene and Aw has three. Muscle 5.1 PPP alignment Jukes-Cantor, neighbor-joining Consensus tree, 1000 bootstrap.

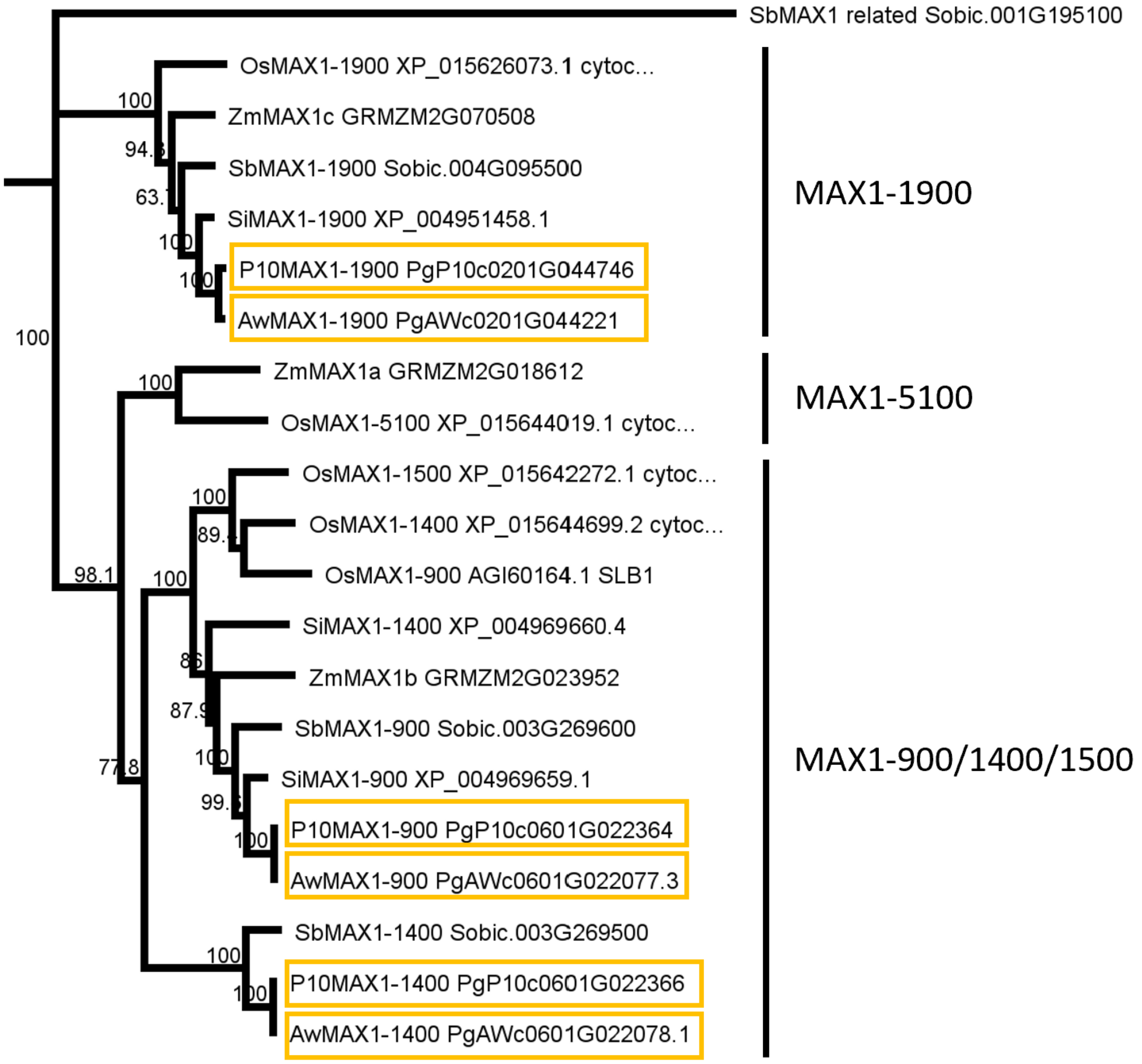

**Supplementary Figure 9**. **Pearl millet MAX1s phylogenetic tree**.

Pearl millet *MAX1* genes (boxed) are named after the closest sorghum homologous gene. The P10 and Aw versions for PgMAX1-900 are identical at the protein level; this is also true for PgMAX1-1400. PgMAX1-900 and PgMAX1-1400 are part of the MAX1-900/1400/1500 subfamily and thus likely use carlactone as a substrate. No clear prediction can be made for the exact enzymatic functions of PgMAX1-900 and PgMAX1-1400 based on this tree, since they don't group cleanly with the OsMAX1s, whose activity has been experimentally confirmed in more detail. Muscle 5.1 PPP alignment Jukes-Cantor, neighbor-joining Consensus tree, 1000 bootstrap; Sobic.001G195100 as outgroup.

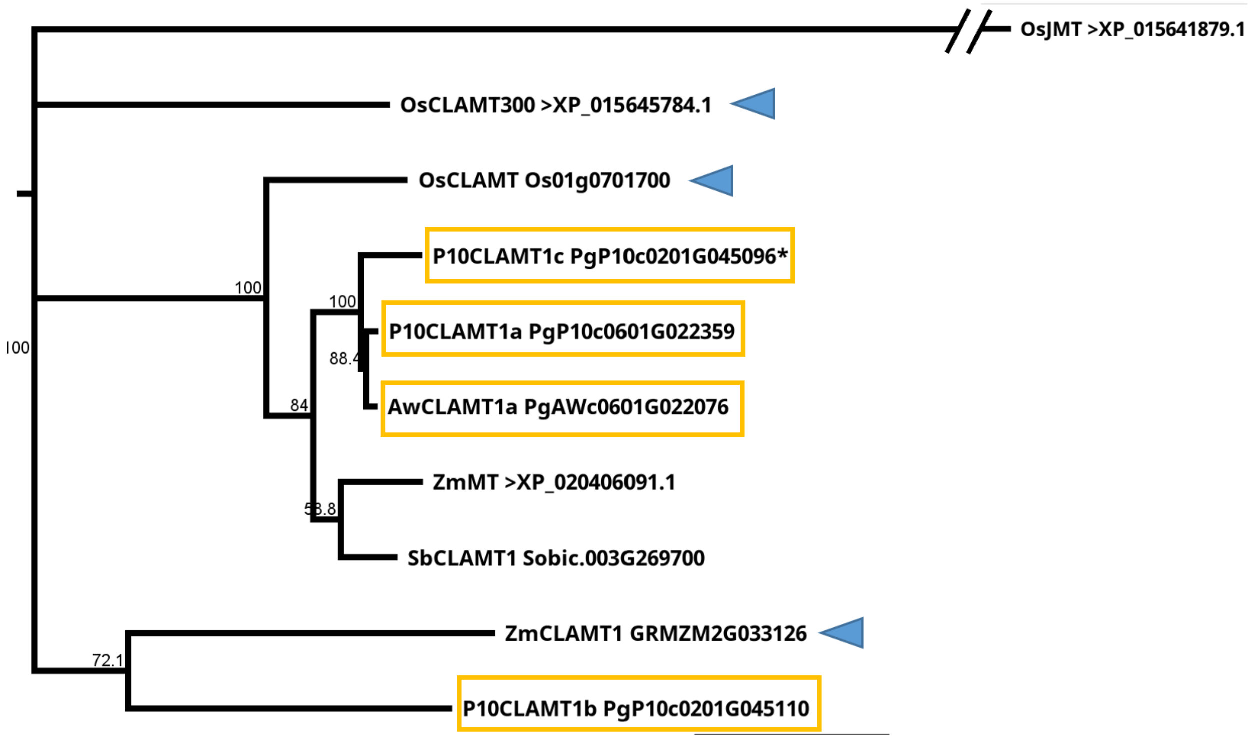

**Supplementary Figure 10**. **Phylogenetic tree of the PgCLAMT gene family**.

There are four PgCLAMTs (boxed); both P10 and Aw have a copy of *CLAMT1a* with a very similar sequence, while P10 has an additional two versions of CLAMT named *P10CLAMT1b* and *P10CLAMT1c*. The *CLAMT* genes with proven enzymatic activity from maize (Li et al., 2023) and rice (Haider et al., 2023) are indicated with arrowheads. Muscle 5.1 PPP alignment Jukes-Cantor, neighbour-joining Consensus tree, 1000 bootstrap; *OsJMT* as outgroup.

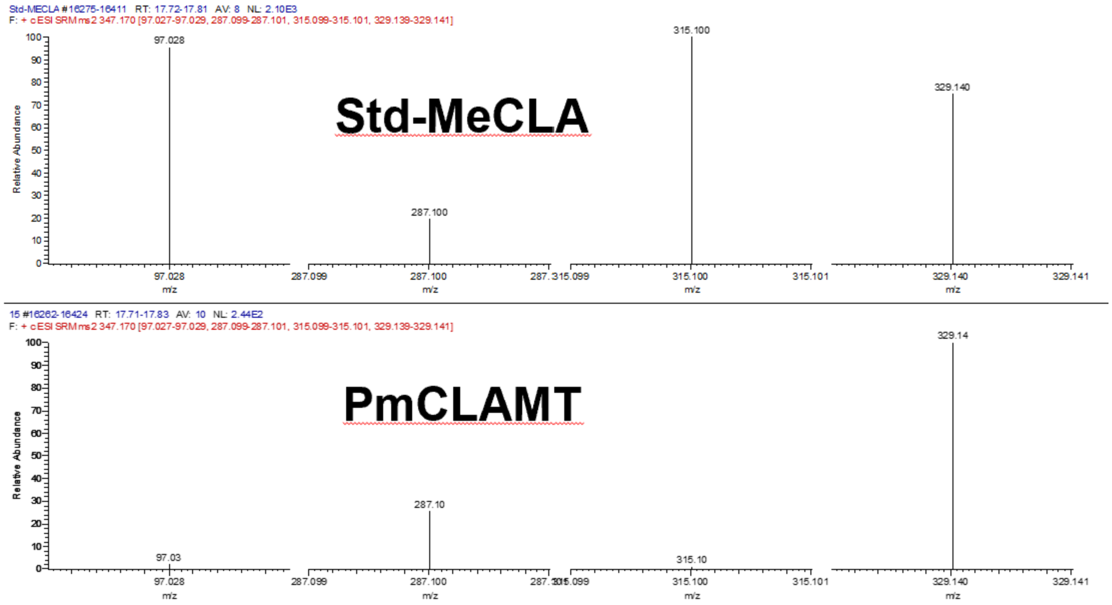

**Supplementary Figure 11. Multiple Reaction Monitoring (MRM) comparison of MeCLA between P10CLAMT1b and MeCLA standard**.

The MRM comparison of the MeCLA standard and the P10CLAMT1b produced MeCLA indicated in Fig1c.

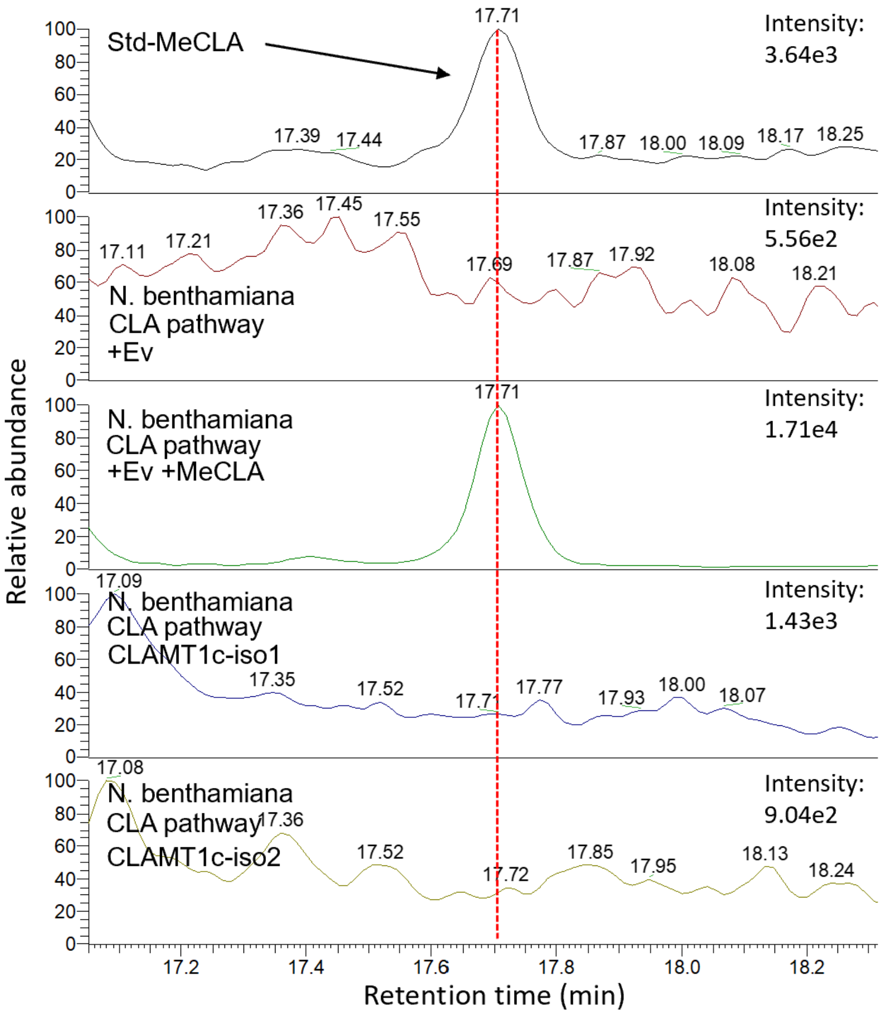

**Supplementary Figure 12. CLAMT1c did not convert CLA into MeCLA by Multiple Reaction Monitoring (MRM) analysis.**

The two P10CLAMT1c predicted isoforms did not convert CLA into MeCLA when tested in the same tobacco leaf transient expression assay as P10CLAMT1b (figure 3).

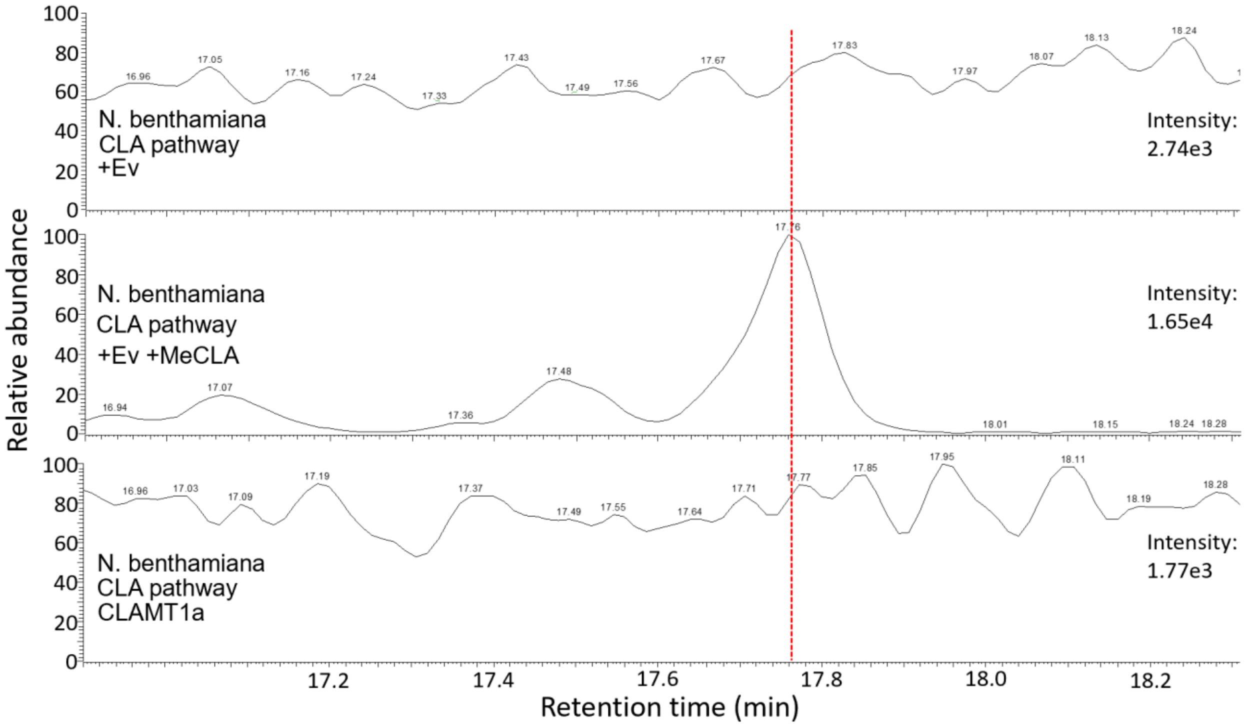

**Supplementary Figure 13. CLAMT1a did not convert CLA into MeCLA by Multiple Reaction Monitoring (MRM) analysis.**

P10CLAMT1a did not convert CLA into MeCLA when tested in the same tobacco leaf transient expression assay as P10CLAMT1b (figure 3).

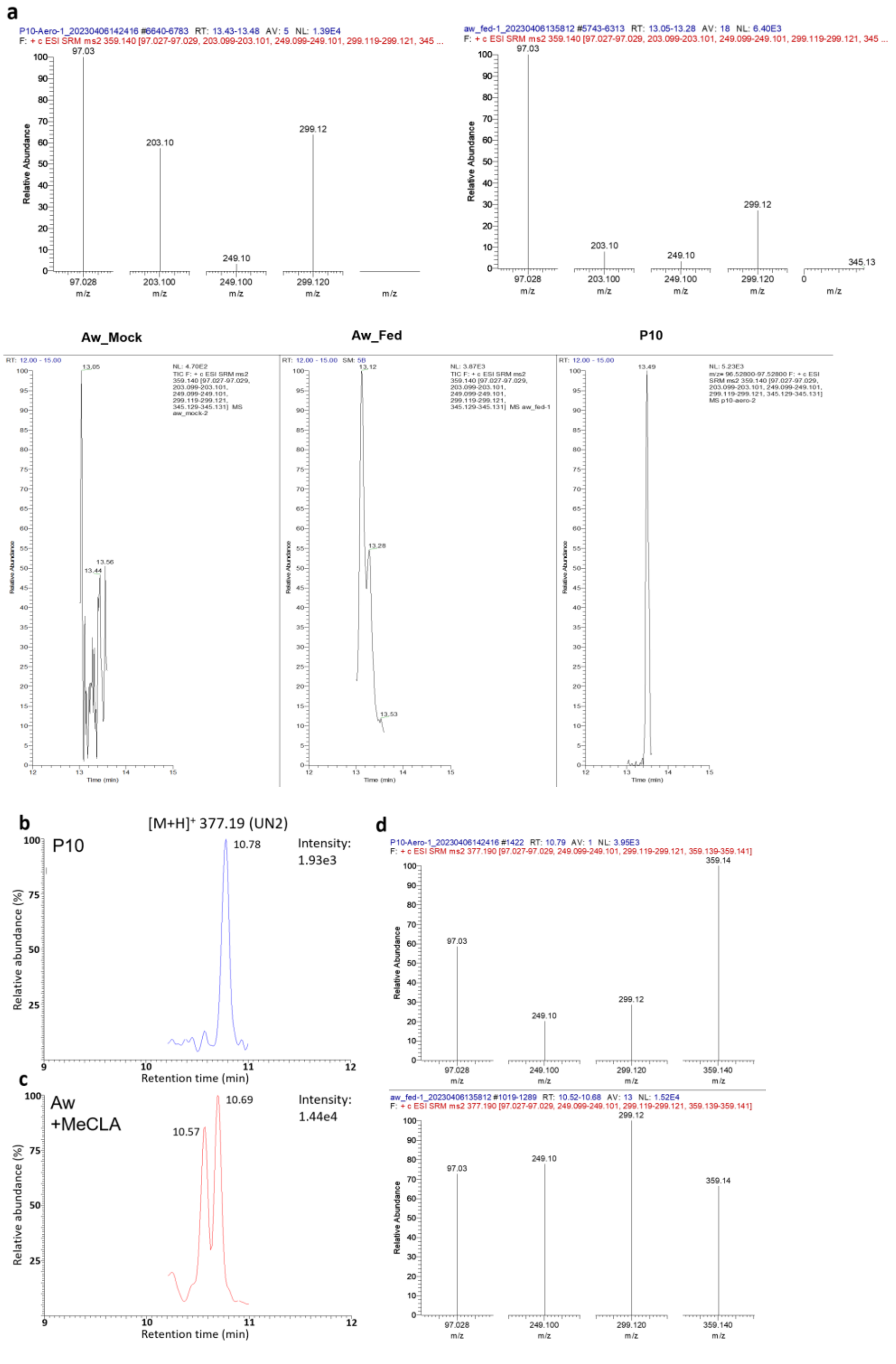

**Supplementary Figure 14. PL2 was produced by Aw, when fed with methyl carlactonoate (MeCLA), by Multiple Reaction Monitoring (MRM) analysis.**

**a** Multiple Reaction Monitoring (MRM) analysis of pennilactone from P10 and the unknown compound of similar retention time produced by Aw when supplied with *rac*-MeCLA showed them to be identical at figure 3c. **b** The new SL PL2 (m/z 377.19) is produced in P10. **c** *rac*-MeCLA-fed Aw also produced a compound at a similar retention time to PL2. **d** Mass fragmentation into the same fragments showed the compound produced by *rac*-MeCLA-fed Aw is identical to P10 PL2.

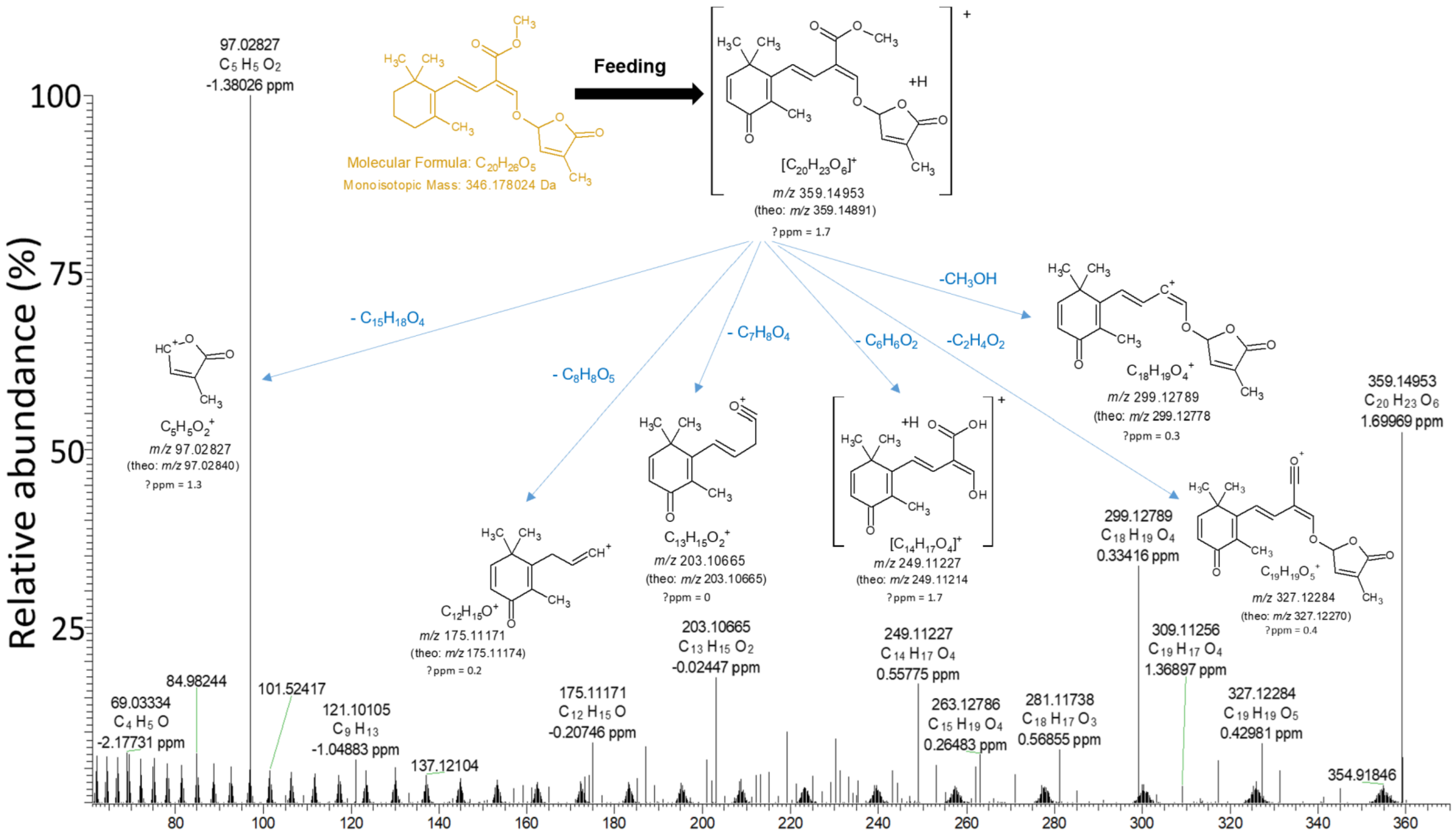

**Supplementary Figure 15. Proposed structure of Pennilactone based on mass fragmentation.**

Proposed structure of Pennilactone (from Pennisetum and strigolactone; $[M+H]^+$ 359.14953 in positive mode) was calculated from MS/MS fragmentation, identified from Aw fed with MeCLA as well as P10 root exudate.

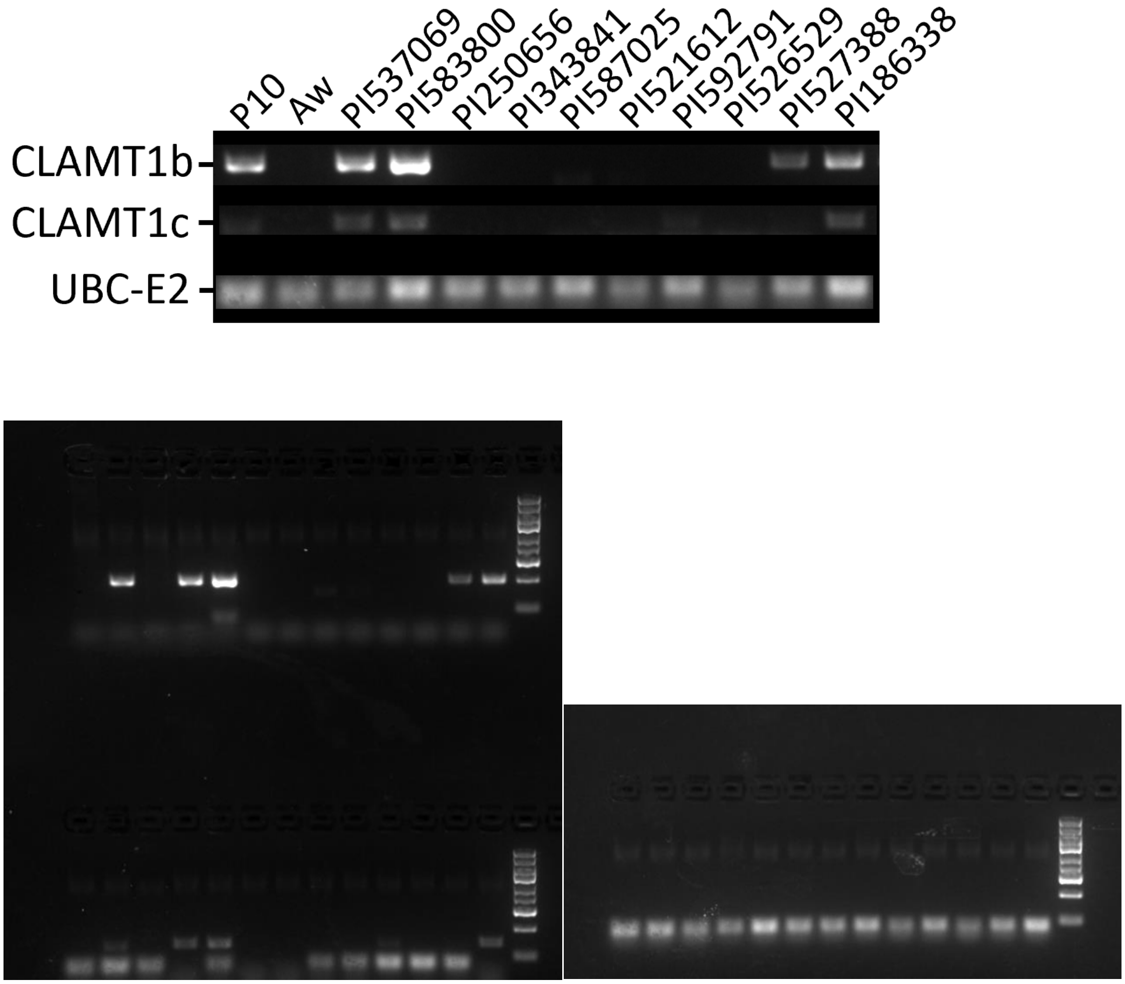

**Supplementary Figure 16. Genotyping 10 accessions for the presence of the *CLAMT1b/c* fragment.**

Genotyping by PCR, for the presence of the *CLAMT1b* and *CLAMT1c* genes, in the 10 lines as received, shows the presence of both genes in P10, PI537069 and PI583800, as expected form their genome sequence. The other eight lines do not show the presence of *CLAMT1b* and *CLAMT1c*, in accordance with their published genomic sequences, except for PI527388 and PI186338. Therefor the PI527388 and PI186338 seeds we received cannot be considered pure lines and were thus excluded from further analysis. In accordance with the guidelines for gel electrophoresis pictures, the unaltered images are provided with the figure.

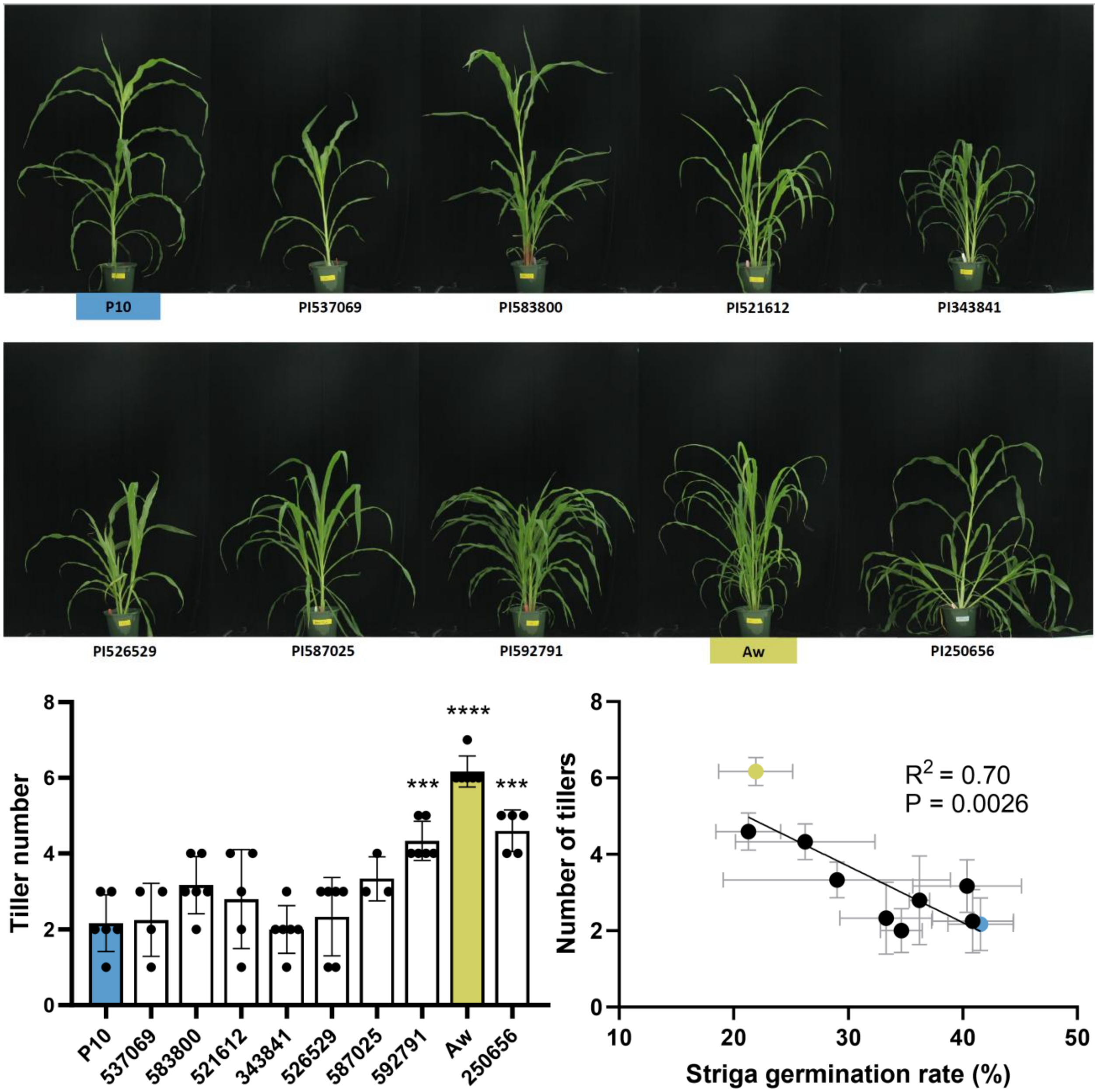

**Supplementary Figure 17. Phenotypes of pearl millet panel accessions.**

**a** The pearl millet panes lines, presented in the order of decreasing induction of *Striga* seed germination by root exudates (Figure 4). The first three lines (P10, PI537069 and PI583800) have the *CLAMT1b* gene and produce pennilactone and other new SLs. **b** Tiller number is only significantly higher than P10 for the three lines with the lowest induction of *Striga* seed germination (PI592791, Aw and PI250656). **c** The correlation between Striga germination rate and number of tillers is negative. With an $R^2$ of 0.70 for a linear regression line with a P-value of 0.0026 the correlation is significant, but not strongly predictive. Therefore, while the PL-branch SLs produced by pearl millet lines likely are affecting tillering, there are additional factors in tiller number determination obscuring a more close correlation.

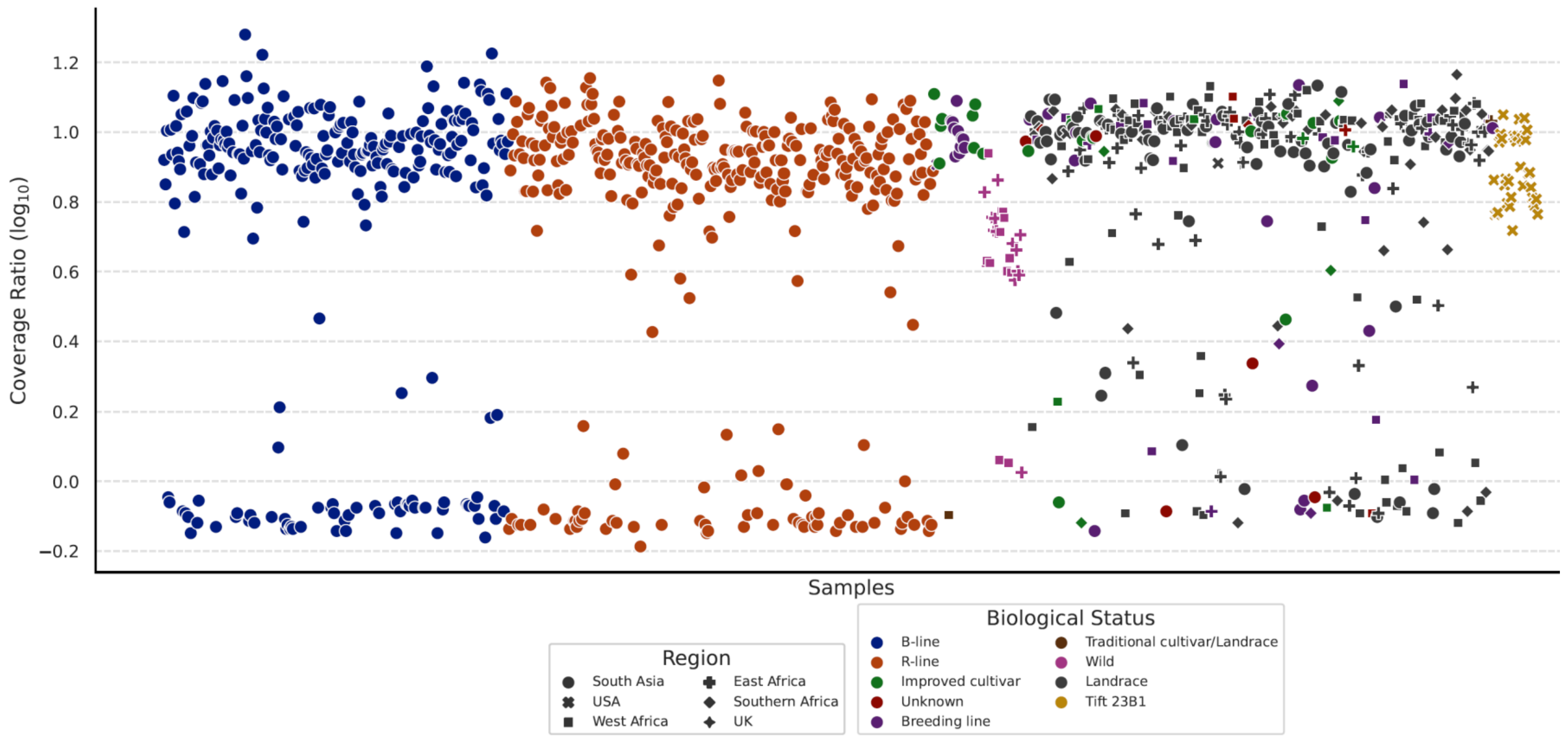

**Supplementary Figure 18**. **Distribution of the CLAMT region in a wide selection of pearl millet accessions.**

Using resequencing data from a wide selection of pearl millet accessions (Varshney et al., 2017), the ratio of the coverage of the P10 chromosome 2 and the coverage of the 0.7Mbp CLAMT region was plotted for each accession. This results in a value of about 1 when the CLAMT region is present, as the coverage is approximately the same as the average coverage of chromosome 2, while the value is much higher when the CLAMT region is absent. The graph shows a clear separation between the two populations and a wide distribution of both accessions with the CLAMT region and without, regardless of provenance. The method of resequencing, RAD or WGS, does not seem to affect the ratio in a significant way. As a control, we included shotgun sequencing data from TIFT-23B1, which does not contain the CLAMT region, and indeed it shows only ratio values above 5.

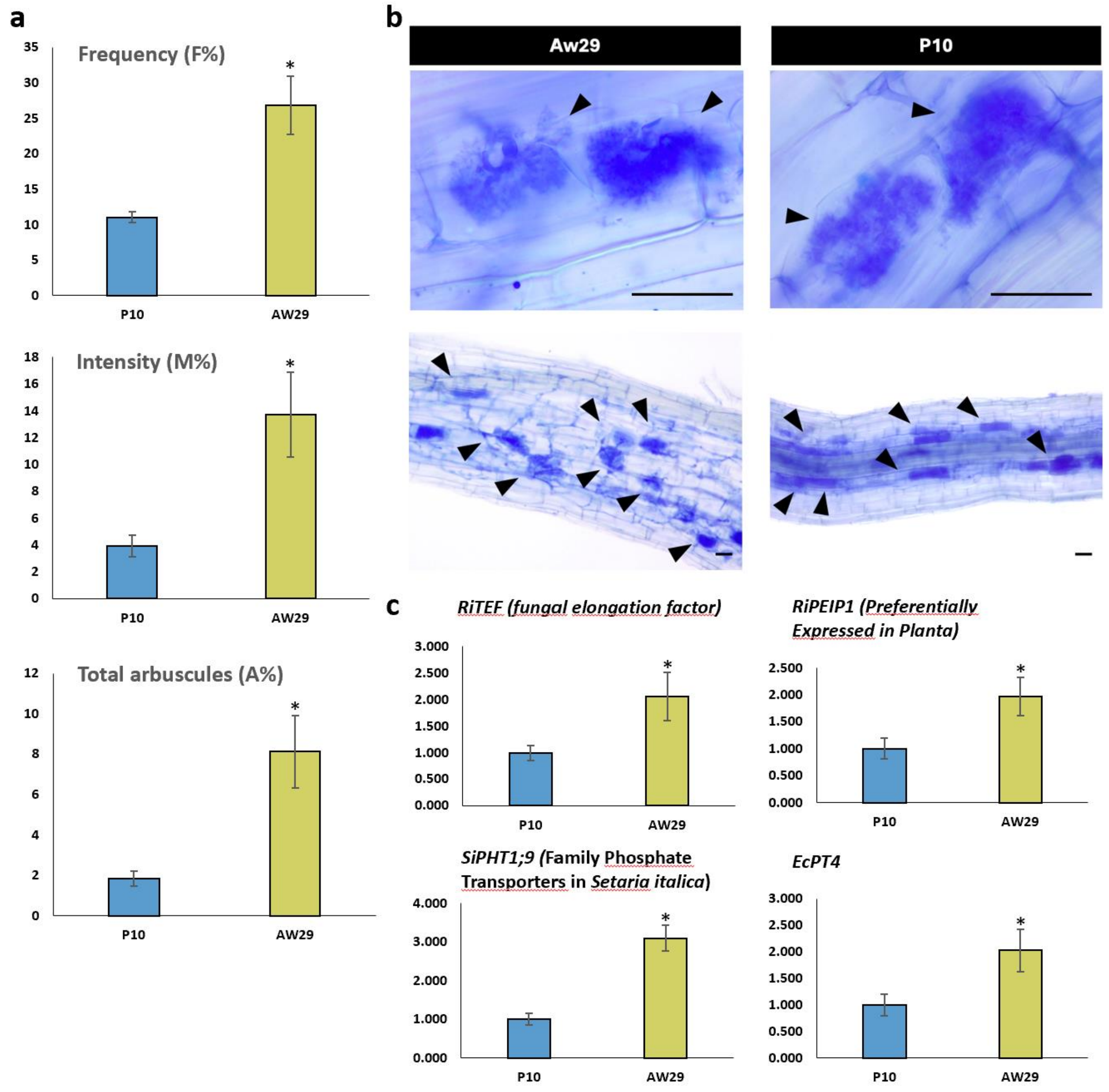

**Supplementary Figure 19. Mycorrhizal colonization of P10 and Aw.**

**a** Morphological evaluation shows that P10 has a significant reduction in both frequency and intensity of colonization compared to Aw. F%: frequency of mycorrhizal colonization; M%: intensity of mycorrhizal colonization; A%: total number of arbuscules. **b** Arbuscule formation at 40 day-post inoculation. Arrows indicate arbuscule-containing cells (Scale bars, 50 $\mu$m). **c** The colonization by the AM fungus *Rhizophagus irregularis* was quantified by measuring the expression of fungal genes (*RiEF* and *RiPEIP1*) and plant AM marker genes (*SiPHT1;9* and *EcPT4*), presented here as normalised expression values. Data are means ± SE (n>=4). Significant values (by One-way ANOVA) are shown as follows: *P < 0.05; **P < 0.01, *** P < 0.001.

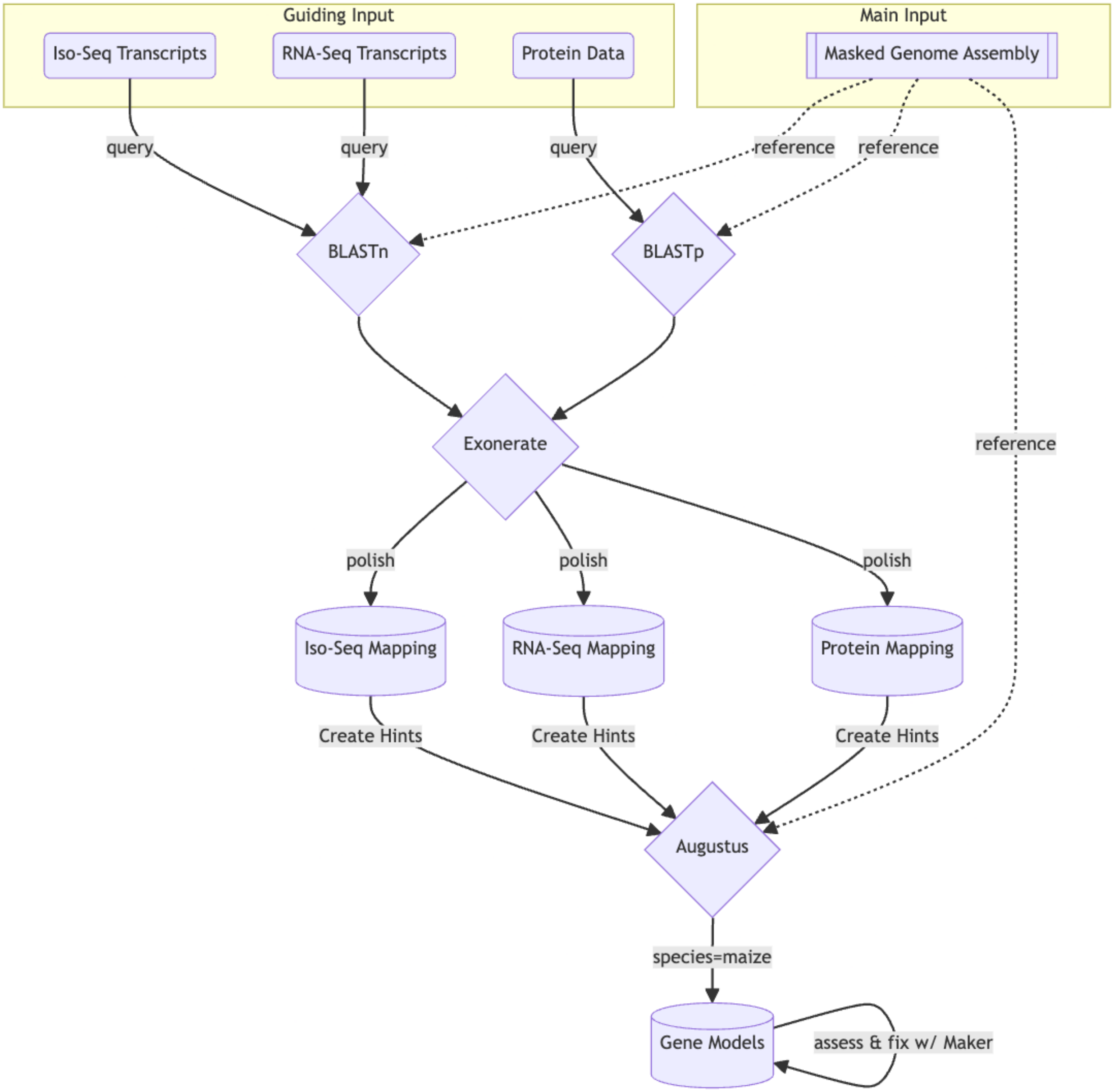

**Supplementary Figure 20. Workflow for Aw and P10 genome annotation.**

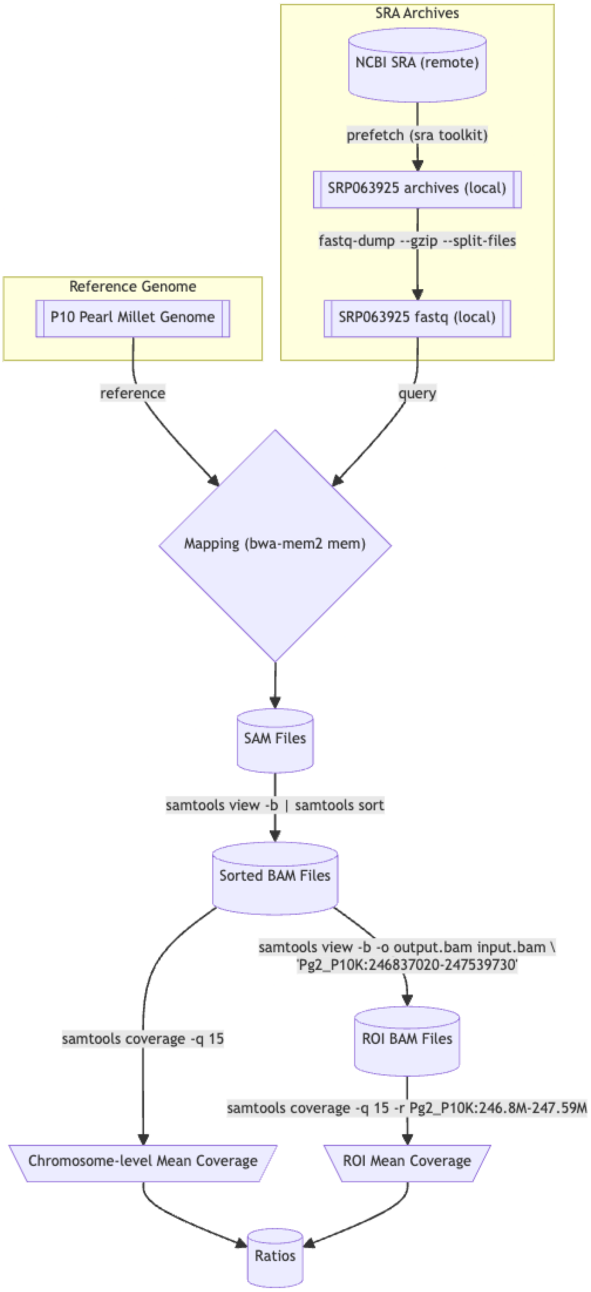

**Supplementary Figure 21. Workflow for analysis of resequencing data.**

**Supplementary Table 1**. **Extended genome assembly and annotation statistics for Aw and P10**.

HC is High Confidence genes and LC is low confidence genes.

[1] Each gap is represented with 100 Ns.

[2] Complete; S: Single BUSCOs; D: Duplicate BUSCOs; F: Fragmented; M: Missing; n: 4896 reference genes from Poales

**Assembly**

| | Aw | | | | P10 | | | |
|---|---|---|---|---|---|---|---|---|
| | *Length* | *GC %* | *GC Skew %* | *Num of gaps* [1] | *Length* | *GC %* | *GC Skew %* | *Num of gaps* [1] |
| Chr1 | 314,009,907 | 48.86 | 0.16 | 0 | 307,615,467 | 48.85 | 0.11 | 0 |
| Chr2 | 276,062,833 | 49.1 | -0.25 | 2 | 281,688,668 | 49.05 | -0.11 | 0 |
| Chr3 | 325,849,051 | 49.21 | 0.09 | 0 | 334,621,887 | 49.22 | -0.01 | 1 |
| Chr4 | 243,773,284 | 48.54 | 0.17 | 2 | 251,295,635 | 48.56 | 0.12 | 1 |
| Chr5 | 174,422,329 | 49 | -0.15 | 1 | 177,041,146 | 49.11 | 0.04 | 0 |
| Chr6 | 286,409,356 | 49.3 | 0.06 | 0 | 282,264,073 | 49.47 | 0.07 | 1 |
| Chr7 | 283,920,974 | 48.96 | 0.02 | 0 | 281,846,746 | 48.91 | -0.03 | 0 |
| ChrUn | 10,392,345 | 53.28 | 1.31 | 152 | 9,861,679 | 47.74 | 1.18 | 235 |

**Annotation**

| | Aw | | | | P10 | | | |
|---|---|---|---|---|---|---|---|---|
| | *Genes/mRNA* | *CDS / Exons* | *5' UTRs* | *3' UTRs* | *Genes/mRNA* | *CDS / Exons* | *5' UTRs* | *3' UTRs* |
| HC | 38,920/62,477 | 284,910/312,473 | 39,586 | 38,211 | 40,689/66,411 | 295,072/320,789 | 38,568 | 37,083 |
| LC | 22,615/25,091 | 44,193/44,253 | 184 | 75 | 23,248/26998 | 49,413/49,470 | 175 | 83 |
| BUSCO [2] | C:93.0%[S:58.7%,D:34.3%],F:2.5%,M:4.5%,n:4896 | | | | C:93.5%[S:59.5%,D:34.0%],F:2.4%,M:4.1%,n:4896 | | | |

**Supplementary Table 2. Repeat content of the Aw and P10 genomes.**

The amount of TE related sequences is highly similar in the two accessions: 1.59 and 1.55 Gbp corresponding to 82.90% and 80.50% of the total genome size for AWK and P10K, respectively. These values are greater than those reported by Varshney et al. in 2017, i.e. 1.22 Gbp and 77.2 % but they entirely meet the expectation Varshney et al made for a true minimum TE fraction of 80% of the entire genome. This testifies the greater efficiency of long read based sequencing techniques in capturing the repetitive fraction of genomes. The most abundant class of TE in Pearl millet is that of Long Terminal Repeat (LTR) retrotransposons totaling to 75.45% and 72.9% in AWK and PK10 genomes, respectively with the Ty3-gypsy superfamily (43.64% and 45.17%) largely more abundant that the Ty1-copia one (12.26% and 17.21%). DNA TEs represent 7.44% and 7.66% of the genome sequence in AWK and PK10, respectively with CACTA and Mutator families being the more abundant (always representing more than 2% of the genome sequence).

| | | Aw | | P10 | |
|---|---|---|---|---|---|
| | | bp | % | bp | % |
| Class 1 | LTR_Copia | 374306143 | 19.55 | 337394745 | 17.52 |
| | LTR_Gypsy | 835647004 | 43.64 | 870042961 | 45.17 |
| | LTR_unknown | 234813472 | 12.26 | 196586386 | 10.21 |
| Class 2 | DNA_HAT | 11691551 | 0.72 | 11672174 | 0.73 |
| | DNA_CACTA | 46761294 | 2.45 | 48386023 | 2.52 |
| | DNA_Harbinger | 15468933 | 0.98 | 16215262 | 0.99 |
| | DNA_Mutator | 26438164 | 2.38 | 25044840 | 2.38 |
| | DNA_Mariner | 15194248 | 0.91 | 16458865 | 0.99 |
| | Total | 1587408978 | 82.90% | 1550644898 | 80.50% |

**Supplementary Table 3**. **List of genes from P10 and Aw which are both highly induced (>2^5) by low phosphate conditions and then induced significantly more by the addition of MP3 treatment**.

The 18 genes in common between P10 and Aw are given a grey background. Homologs of enzymes reported to be part of the strigolactone biosynthesis pathway in maize are highlighted in yellow. In the case of *CCD8*, *CLAMT* and *MAX1* these are not the only homologous options in pearl millet (see supplemental figures 8, 9 and 10), but we can conclude from this co-expression that these are the most likely candidates to be active genes in the strigolactone biosynthetic pathway. The candidate genes for the production of Pennilactone and PL2 from methyl carlactonoate can be upregulated in both P10 and Aw, however for the genes involved in producing PL3 and PL4 this is not expected, since these were not detected after feeding Aw with methyl carlactonoate.

| P10 lowP/MP3 up | description from similar proteins | Aw lowP/MP3 up |
|---|---|---|
| PgP10c0101G010112 | Tropinone reductase | PgAWc0101G010023 |
| | Obtusifoliol demethylase | PgAWc0101G011320 |
| | Chalcone synthase | PgAWc0101G011727 |
| PgP10c0101G011332 | D27 | |
| PgP10c0101G012222 | probable carboxylesterase | |
| PgP10c0101G012240 | Abscisic acid 8'-hydoxylase 3 | |
| PgP10c0101G017802 | benzyl alcohol O-benzoyltransferase | PgAWc0101G017579 |
| | hypothetical protein | PgAWc0201G036589 |
| | F-box/FBD/LRR-repeat protein At1g13570-like | PgAWc0201G036590 |
| | Ergosterol biosynthetic protein 28 | PgAWc0201G038150 |
| | cytochrome P450 89A2 | PgAWc0201G038988 |
| PgP10c0201G040400 | ent-copalyl diphosphate synthase 1, chloroplastic | PgAWc0201G039931 |
| | protein CYCLOPS-like | PgAWc0201G040446 |
| PgP10c0201G042842 | putative 2-oxoglutarate-dependent dioxygenase | PgAWc0201G042298 |
| PgP10c0201G042843 | ent-isokaurene C2-hydroxylase | PgAWc0201G042299 |
| | unknown | PgAWc0201G044004 |
| | unknown | PgAWc0201G044005 |
| PgP10c0201G045110 | CLAMT1b | |
| | bidirectional sugar transporter SWEET15 | PgAWc0301G003590 |
| | UDP-glycosyltransferase 85A2 | PgAWc0301G005124 |
| PgP10c0301G006912 | ABC transporter B family member 15 | PgAWc0301G006836 |
| | hypothetical protein | PgAWc0401G047069 |
| | S-norcoclaurine synthase 1 | PgAWc0401G047821 |
| PgP10c0401G048472 | 2-oxoglutarate-dependent dioxygenase - S-norcoclaurine synthase 1 | |
| PgP10c0401G048625 | 2-oxoglutarate-dependent dioxygenase - S-norcoclaurine synthase 1 | |
| PgP10c0401G048754 | indole-2-monooxygenase | PgAWc0401G048099 |
| PgP10c0401G049555 | aquaporin NIP3-3 | |
| PgP10c0401G051155 | CYP706 | PgAWc0401G050240 |
| PgP10c0401G051273 | uncharacterized protein | |
| | 2-alkenal reductase like | PgAWc0401G052246 |
| PgP10c0401G053327 | ent-copalyl diphosphate synthase 2 | PgAWc0401G052458 |
| PgP10c0501G055622 | NADP-dependent alkenal double bond reductase P2-like | |
| PgP10c0501G055754 | cytochrome P450 93A3 | |
| PgP10c0501G056021 | cytochrome P450 93A3 | |
| | squamosa promoter-binding-like | PgAWc0501G055634 |
| PgP10c0501G058210 | serine carboxypeptidase-like 34 | PgAWc0501G057204 |
| | cytochrome P450 76M5 | PgAWc0501G058053 |
| | serine/threonine receptor-like kinase NFP | PgAWc0501G059262 |
| PgP10c0601G019773 | Tropinone reductase-like protein | |
| PgP10c0601G021829 | CCD8 | PgAWc0601G021537 |
| PgP10c0601G022089 | ABC transporter G family member 38-like | PgAWc0601G021810 |
| PgP10c0601G022322 | germin-like protein 1-2 | PgAWc0601G022036 |
| PgP10c0601G022366 | MAX1-1400 | PgAWc0601G022078 |
| PgP10c0601G023133 | putative fatty-acid--CoA ligase | PgAWc0601G022896 |
| PgP10c0601G027441 | hypothetical protein | |
| | short-chain dehydrogenase TIC 32, chloroplastic | PgAWc0701G027593 |
| | ABC transporter B family member 4 | PgAWc0701G027594 |
| | cytochrome P450 709B2 | PgAWc0701G028056 |
| PgP10c0701G029515 | cytochrome P450 716B1-like | PgAWc0701G029274 |
| PgP10c0701G029516 | cytochrome P450 716B1 | PgAWc0701G029275 |
| PgP10c0701G031847 | putative amidase At4g34880-like | PgAWc0701G031529 |
| | unknown | PgAWc0701G032050 |
| | ZAS | PgAWc0701G032297 |
| | epoxide hydrolase A-like | PgAWc0701G034036 |
| | putative 1-deoxy-D-xylulose-5-phosphate synthase 2, chloroplastic | PgAWc0701G034758 |

**Supplementary Table 4**. **Presence (green) or absence (red) of genes in the *CLAMT* gap and flanking areas of chromosome 2 for P10, Aw, and the 10 sequenced pearl millet lines described by Yan et al. (2023).** Numbers in each cell indicate the percentage of the protein sequence that is identical to P10, showing that the CLAMT1b proteins produced by P1537069 and P1583800 are identical to the P10 CLAMT1b.

| | | flavonoid methyltra | WVD2-like | CLAMT1c | acyl trans-ferase | CYP51 | CLAMT1b | Glucosi-dase | SEC14 |
|---|---|---|---|---|---|---|---|---|---|
| P10 | | | | | | | | | |
| Aw | | | | | | | | | |
| PI592791 | Tifleaf 3 | 98 | 95 | | | | | 96 | 97 |
| PI537069 | Baoudarache | 99 | 99 | 99.94 | 99.77 | 99.90 | 100 | 100 | 99 |
| PI583800 | ICMV 88908 | 99 | 99 | 99.26 | 99.77 | 99.93 | 100 | 96 | 99 |
| PI343841 | - | 99 | 98 | | | | | 97 | 99 |
| PI527388 | 11 | 98 | 96 | | | | | 95 | 97 |
| PI526529 | AMM 1227 | 98 | 96 | | | | | 96 | 97 |
| PI521612 | Ngululu | 100 | 99 | | | | | 96 | 97 |
| PI587025 | Dokhn | 98 | 96 | | | | | 97 | 97 |
| PI250656 | K551 | 98 | 96 | | | | | 95 | 97 |
| PI186338 | - | 98 | 96 | | | | | 95 | 97 |

**Supplementary Table 5. Candidate genes for affecting striga resistance as reported from GWAS experiments in sorghum (Mallu et al., 2022; Kavuluko et al 2021) and maize (Badu-Apraku et al., 2020) with the corresponding homologous genes identified in P10 and Aw**.

The difference in amino acids between the predicted proteins from P10 and Aw are listed, with the two noted outliers MAKR3 and WRKY30 highlighted.

We took the genes with significant association to Striga susceptibility from these studies and compared the sequence differences between the homologous genes in P10 and Aw. Of the 65 gene pairs analyzed 63 have fewer than 10 amino acids difference between P10 and Aw, with a total average of 4.02 AA and a median of 2 AA difference (supplemental table 5). Clear outliers PgMAKR3 (PgP10c0401G048596.1 / PgAWc0401G047945 + PgAWc0401G047946) show 102 AA and PgWRKY30 (PgP10c0701G036602.1 / PgAWc0701G036159) 24 AA difference between P10 and Aw. In contrast to its annotation, PgMAKR3 is a CC-NB-LRR resistance gene (Supplemental figure 6) and most of the differences between the P10 and Aw gene are located in the variable LRR domain, which is involved in pathogen recognition. Pathogen recognition by CC-NB-LRR proteins is translated into an immune response by downstream WRKY transcription factors, a pathway that has been suggested for resistance to parasitic plants (Hu et al., 2020). PgWRKY30 could be the WRKY TF paired with PgMAKR3 in the recognition of a pathogen, such as *Striga*. This finding could be a starting point towards discovering more about the differences in post-attachment resistance between Aw and P10.

| name | sp. | code | long name/function | Source | P10 geneID | Aw geneID | AA diff |
|---|---|---|---|---|---|---|---|
| ABI5 | Sb | Sobic.003G363400 | Abscisic acid-insensitive 5 (ABI5) | Mallu 2022 | PgP10c0601G020578.2 | PgAWc0601G020282 | 5 |
| NINJA | Sb | Sobic.001G335400 | Novel interactor of Jaz (Ninja) | Mallu 2022 | PgP10c0501G058583.1 | PgAWc0501G057612 | 1 |
| MAKR3 | Sb | Sobic.009G020600 | Membrane-associated kinase regulator 3 | Mallu 2022 | PgP10c0401G048596.1 | PgAWc0401G047945 + PgAWc0401G047946 | 102 |
| Masp | Sb | Sobic.001G263200 | α/β hydrolase/Maspardin | Mallu 2022 | PgP10c0201G040775.3 | PgAWc0201G040281 | 0 |
| AGP | Sb | Sobic.002G047700 | ADP-glucose phosphorylase | Mallu 2022 | PgP10c0601G021848.1 | PgAWc0601G021565.2 | 2 |
| AGPa | | | | | PgP10c0101G017229.2 | PgAWc0101G017022.2 | 1 |
| | Sb | Sobic.003G246200 | unknown | Mallu 2022 | PgP10c0601G022818.4 | PgAWc0601G022552.3 | 1 |
| DOR | Sb | Sobic.004G050300 | Disulphide oxido reductase/ uncharacterized | Mallu 2022 | PgP10c0401G046705.1 | PgAWc0401G046094.2 | 2 |
| | Sb | Sobic.008G025300 | unknown | Mallu 2022 | PgP10c0101G019321.1 | PgAWc0101G019027 | 7 |
| LGS1/CST! | Sb | Sobic.005G213600 | cytosolic sulfotransferase 5 | Mallu 2021 | PgP10c0101G011829.1 | PgAWc0101G011593.4 | 0 |
| | | | | | PgP10c0101G010309.1 | PgAWc0101G010195.1 | 2 |
| | | | | | PgP10c0201G042252.2 | PgAWc0201G041717.2 | |
| ABC-G | Sb | Sobic.002G321300 | pleiotropic drug resistance (PDR)/ ATP Bindin | Kavuluko 2021 | PgP10c0701G029502.1 | PgAWc0701G029263.2 | 5 |
| IFR | Sb | Sobic.003G241300 | Isoflavon reductase | Kavuluko 2021 | PgP10c0601G025943.1 | PgAWc0601G025783.x | 1 |
| | | | | | PgP10c0601G025942.1 | PgAWc0601G025783.1 | 1 |
| FLAP11 | Sb | Sobic.009G056400 | Fasciclin-like arabinogalactan protein 11 | Kavuluko 2021 | PgP10c0101G017488.1 | PgAWc0101G017280 | 1 |
| PMT2 | Sb | Sobic.002G201600 | Methyltransferase | Kavuluko 2021 | PgP10c0701G031772.2 | PgAWc0701G031602.1 | 0 |
| NAC4 | Sb | n/a | Secondary wall NAC transcription factor 4 | Kavuluko 2021 | PgP10c0401G048746.1 | PgAWc0401G048090 | 0 |
| ENOD93 | Sb | Sobic.010G032000 | Early nodulin 93 | Kavuluko 2021 | PgP10c0201G041260.1 | PgAWc0201G040753 | 2 |
| | | | | | PgP10c0201G041265.1 | PgAWc0201G040760 | 6 |
| Xin1 | Sb | Sobic.005G099000 | Xylanase inhibitor 1 | Kavuluko 2021 | PgP10c0401G052759.1 | PgAWc0401G051935 | 2 |
| ERF113 | Sb | Sobic.004G158901 | Ethylene response factor | Kavuluko 2021 | PgP10c0301G003555.1 | PgAWc0301G003515.1 | 3 |
| Paco1 | Sb | Sobic.010G049100 | Peroxisomal acyl-CoA oxidase 1 | Kavuluko 2021 | PgP10c0201G041611.1 | PgAWc0201G041104.x | 0 |
| DMR6 | Sb | Sobic.006G190000 | Downy mildew resistance 6 | Kavuluko 2021 | PgP10c0501G061435.1 | PgAWc0501G060409 | 0 |
| PRX1 | Sb | Sobic.006G009400 | Peroxiredoxin1 | Kavuluko 2021 | PgP10c0501G054714.1 | PgAWc0501G053791.1 | 2 |
| | Sb | Sobic.004G005100 | zinc finger with peptidase domain | Kavuluko 2021 | PgP10c0201G046565.1 | PgAWc0201G045955 | 0 |
| HSP60 | Sb | Sobic.007G004500 | Heat shock protein 60 | Kavuluko 2021 | PgP10c0401G048761.3 | PgAWc0401G048105.3 | 2 |
| WRKY30 | Zm | GRMZM2G143204 | wrky30—WRKY-transcription factor 30 | Badu-Apraku 2020 | PgP10c0701G036602.1 | PgAWc0701G036159 | 24 |
| EREB184 | Zm | GRMZM2G028151 | ereb184—AP2-EREBP-transcription factor 18 | Badu-Apraku 2020 | PgP10c0201G037684.1 | PgAWc0201G037222 | 2 |
| MYB51 | Zm | GRMZM5G803355 | myb51—MYB-transcription factor 51 | Badu-Apraku 2020 | PgP10c0101G018838.1 | PgAWc0101G018548.1 | 0 |
| MYB8 | Zm | GRMZM2G041415 | myb8—myb transcription factor8 | Badu-Apraku 2020 | PgP10c0401G051138.1 | PgAWc0401G050228.1 | 0 |
| | | | | | PgP10c0701G030347.2 | PgAWc0701G030079.3 | 0 |
| bZIP8 | Zm | GRMZM2G146020 | bzip8—bZIP-transcription factor 8 | Badu-Apraku 2020 | PgP10c0101G019027.1 | PgAWc0101G018740.2 | 3 |
| PSK2 | Zm | GRMZM2G079290 | psk2—phytosulfokine2 | Badu-Apraku 2020 | PgP10c0401G054113.1 | PgAWc0401G053145.1 | 0 |
| | | | | | PgP10c0401G054111.1 | PgAWc0401G053143.1 | 0 |
| bHLH150 | Zm | GRMZM2G045431 | bhlh150—bHLH-transcription factor 150 | Badu-Apraku 2020 | PgP10c0701G031420.1 | PgAWc0701G031115.1 | 0 |
| HSFTF21 | Zm | GRMZM2G139535 | hsftf21—Heat shock factor protein 4 | Badu-Apraku 2020 | PgP10c0701G031397.1 | PgAWc0701G031097.1 | 0 |
| bHLH105 | Zm | GRMZM2G082586 | bhlh105—bHLH-transcription factor 105 | Badu-Apraku 2020 | PgP10c0701G031182.1 | PgAWc0701G030876.1 | 1 |
| EREB71 | Zm | GRMZM2G113060 | ereb71—AP2-EREBP-transcription factor 71 | Badu-Apraku 2020 | PgP10c0501G060646.1 | PgAWc0501G059638.1 | 3 |
| bHLH97 | Zm | AC149829.2_FG00 | bhlh97—bHLH-transcription factor 97 | Badu-Apraku 2020 | PgP10c0501G060900.1 | PgAWc0501G059901.1 | 0 |
| HSFTF20 | Zm | GRMZM2G301485 | hsftf20—HSF-transcription factor 20 | Badu-Apraku 2020 | PgP10c0301G008054.1 | PgAWc0301G008012.1 | 6 |
| bHLH55 | Zm | GRMZM2G030762 | bhlh55—putative DNA-binding domain super | Badu-Apraku 2020 | PgP10c0601G019990.2 | PgAWc0601G019678.2 | 3 |
| MGT8 | Zm | GRMZM2G065971 | mgt8—magnesium transporter8 | Badu-Apraku 2020 | PgP10c0601G020072.1 | PgAWc0601G019759.1 | 3 |
| CYP26 | Zm | GRMZM2G087875 | cyp26—putative cytochrome P450 superfamil | Badu-Apraku 2020 | PgP10c0601G020613.1 | PgAWc0601G020312.1 | 1 |
| MYB79 | Zm | GRMZM2G056986 | MYB-type transcription factor 79 | Badu-Apraku 2020 | PgP10c0601G020644.1 | PgAWc0601G020341.1 | 3 |
| | | | | | PgP10c0101G015419.1 | PgAWc0101G015249.1 | 1 |
| HCF60 | Zm | GRMZM2G038013 | hcf60—high chlorophyll fluorescence60 | Badu-Apraku 2020 | PgP10c0301G009454.2 | PgAWc0301G009405.2 | 9 |
| | | | | | PgP10c0301G009482.2 | PgAWc0301G009430.1 | 3 |
| MYB | Zm | GRMZM2G169316 | MYB transcription factor | Badu-Apraku 2020 | PgP10c0601G020688.1 | PgAWc0601G020389.1 | 8 |
| HSP1 | Zm | GRMZM2G310431 | hsp1—heat shock protein | Badu-Apraku 2020 | PgP10c0601G020804.1 | PgAWc0601G020534.1 | 2 |
| | | | | | PgP10c0101G015535.1 | PgAWc0101G015357.1 | 0 |
| | | | | | PgP10c0101G009566.1 | PgAWc0101G009510.1 | 7 |
| | | | | | PgP10c0501G059814.1 | PgAWc0501G058825.1 | 0 |
| | | | | | PgP10c0401G047662.1 | PgAWc0401G047029.1 | 3 |
| NACtf20 | Zm | GRMZM2G180328 | nactf20—NAC-transcription factor 20 | Badu-Apraku 2020 | PgP10c0101G015152.1 | PgAWc0101G014996.4 | 2 |
| NACtf112 | Zm | GRMZM2G456568 | nactf112—NAC-transcription factor 112 | Badu-Apraku 2020 | PgP10c0101G015184.2 | PgAWc0101G015019.3 | 5 |
| MYBr3 | Zm | GRMZM2G002128 | mybr3—MYB-related-transcription factor 3 | Badu-Apraku 2020 | PgP10c0101G015419.1 | PgAWc0101G015249.1 | 1 |

**Supplementary Table 6. Gene sequences used in this study**

| Gene | Coding sequence |
|---|---|
| **CLAMT1a** | ATGGCGTCCTCGCTGCTCCACTGCTCCGACAAGCTCCCGTTCATGGACGTGGAGACAATCCTCCACATGAAAGAGG<br>GGCTTGGCGAGACCAGCTACGCGCAGAACTCCTCTCTTCAGAAGCGGGGCATGGACACGCTGAAGAGCCTCATCAC<br>CAACTCGGCGACGGACGTGTACATCTCGCAGATGCCGGAGAGGTTCACGGTGGCCGACCTGGGCTGCTCGTCGGGC<br>CCGAACGCACTGTGCCTCGTCGAGGACATCGTCGGGAGCATCGGCCGGGTGTGCGGCCGGTCGTCGCAGCCGCCGC<br>CCGAGTTCTCGGTGCTCCTCAACGACCTCCCGACCAACGACTTCAACACCATCTTCTTCAGCCTGCCGGAGTTCAC<br>CGACCGGCTCAAGGCCGCCGCCGAGACCGACGAGTGGGGCCGGCCGATGGTGTTCCTGTCCGGCGTGCCGGGGTCT<br>TTCTACGGGAGGCTCTTCCCCAGGAAGAGCGTGCACTTCATCTGCTCCTGCTCCAGCCTGCACTGGCTCTCCCAGG<br>TCCCGCCGGGGCTCTTCGACGAGGCCACGGGCACGCCCATCAACAAGGGGAAGATGTACATCTCGAGCTCCAGCCC<br>GCTCGCCGTGCCGACGGCCTACCTGAGGCAGTTCCAGAGGGACTTCGGCCTGTTCCTCAGATCGCGCGCCGCCGAG<br>GTCGTCGCCGGCGGCCGGATGGTACTGGCCATGCTCGGCAGGCAGACCGAGGGCTACATCGACAGGCGAACCACCT<br>TCCTCTGGGAGCTCCTCTCCGAGTCGTTCGCCTCGCTCGTGGCACAGGGGTTGGTGGCCCAGGAGAAGGTGGACGC<br>GTACAACGTGCCGTTCTACGCGCCGTCGATCCAGGAGGTGGAGGAGGAGGTGCGGCGAGAGGGGTCGTTCCGGCTC<br>GACTACGTGCAGACGTACGAGATCAACCTGAGCAGCAGCGGTGACGCCAAGGAGGACGGCCGGACGGTGTCCATGG<br>CGATCAGGGCCATCCAGGAGTCCATGCTGAACCACCACTTCGGCCCAGACATTGTCGACGCGCTCTTTCACAGGTA<br>CACGGAACTCGTCACCGAGTCCATGGAGAGGGAGGAGGTCGAAAGCGTTCAGATTGGGGTCGTCCTCACAAGGTTA<br>TGA |
| **CLAMT1b** | ATGGCTTCTTCCTCCAATTGCTCCAACAAACACCCTCCCATGATGAATGTGGAGGCCGTCCTCCACATGAAGGCAG<br>GCCTCGGTGAGAACAGTTACGCGATGAACTCTGTGATTCAGAAAAAAGGCATGGACACCGTGAGGAGCCTTGTCAC<br>CGATTCAGCGACACAACTGTACTTGTCACTCAAACCAGAGAGGTTCACCATGGCCGACCTTGGGTGCTCCTCGGGG<br>ACGAACGCGCTGGGCATGGTGGAGGACGTCGTCAGGAGCGTCGGCAAGGCCTGCTGTGGCTGCGGGTCACCGCCGC<br>CAGAGTTCTCGGTGCTCCTGAACGATCTCCCTACCAACGACTTCAACACCGTCTTCTCCCGGGTACCAGAGTTCAC<br>AGCCAAGCTGAAAGCCGACGCCCCGTCGCCGATGATCTTCGTCTCTGGCGTCCCGGGATCCTTCTACGGAAGGCTG<br>TTCCCTAGCAGGAGCGTGCACTTGGCTTGCTCCTTCGCTAGCCTGCACTGGCTCTCTCAGGTTCCTCGACTTTCTG<br>ATGAGACGAACACGCCCATGAACAAGGGGAAGATGTTCATATCAAGCACGAGCCCTCCTGCCGTGCCGGCGGCCTA<br>CCTGAGGCAGTTCCAGAGAGACTTCACCGTTTTCCTCGAGTCACGAGCCGCGGAGGTTGTCCCCGGTGGCCGCATA<br>GTTGTGTCGATGCTAGGGCGTCAAACCGAGGGGTACACGGACATGAAGACCACCCTCTTGTGGGATCTTCTTTCCG<br>AGTCGCTTGCCGCGATGGTTTCACAGGGCTTGGCCGAGCAGGAGAAGGTCGATGCCTACGACGTTCCCTTCTACGC<br>GCCAAGCTTACGGGAGATCAAGGAAGTGGTGAGCAAGGAAGGTTCCTTCAGCCTCAACTGCGTGAGGACGTACGAG<br>GCAACCCATGGCGGGAGCGACGCCAAAAGAGACGGCAAGATGCTAGCTATGACGGCCAGGGCCGTCCATGAGTCGA<br>TGATGAGCCACCACTTCGGGCCAGGCATCGTGGAGCCGCTGTTCCACAAGTACGGCGAGTTGGTTGCCCAGTTCAT<br>GGAGACGGGAGAGGTCAAGAGTGTCCAGATAGGGGTGGTCCTGACAAGGTTGTTGTGA |
| **CLAMT1c<br>-iso1** | ATGGCGTCCTCGCTGCTCCACTGCTCCGACAAGCTCCCGTTCATGGACGTGGAGACCATCCTCCACATGAAAGAGG<br>GGCTCGGCGAGACCAGCTACGCGCAGAACTCCTCGCTTCAGAAGCGGGGCATGGACACGCTGAAGAGCCTTATCAC<br>CAACTCGGCGACGGACGTGTACATCGCGCAGATGCCGGAGAGGTTCACGGTGGCCGACCTGGGCTGCTCGTCGGGC<br>CCGAACGCGCTGTGCCTCGTCGAGGACATCGTCGGGAGCATCGGCCGGGTGTGCGGCCGGTCGTCGCAGCCGCCGC<br>CCGAGTTCTCGGTGCTCCTCAACGACCTCCCGACCAACGACTTCAATACCATCTTTTTCAGCCTGCCGGAGTTCAC<br>CGACCGGCTCAAGGCCGCCGCCGAGACCGATGAGTGGGGCCGGCCGATGGTGTTCCTGTCCGGCGTGCCGGGGTCC<br>TTCTACGGGAGGCTCTTCCCAAGGAAGAGCGTGCACTTCATCTGCTCCTACTCCAGCTTGCACTGGCTCTCCCAGG<br>TCCCGCCGGGGCTCTTCGACGAGGCCACGGGCACGCCCATCAACAAGGGGAAGATGTACATCTCGAGCTCCAGCCC<br>GCTCGCCGTGCCGACAGCCTACCTGAGGCAGTTCCAGAGGGACTTCGGCCTGTTCCTCAGATCACGCGCCGCCGAG<br>GTCGTCGCCGGCGGCCGGATGGTCCTCGCCATGCTCGGCAGGCAGACCGAGGGCTACATCGACAGGCGAACCACCT<br>TCCTCTGGGAGCTCCTCTCCGAGTCGTTCGCCTCGCTCGTGGCACAGAACGCCGAAGAACCTGAACTTGATTACTG<br>A |
| **CLAMT1c<br>-iso2** | ATGGCGTCCTCGCTGCTCCACTGCTCCGACAAGCTCCCGTTCATGGACGTGGAGACCATCCTCCACATGAAAGAGG<br>GGCTCGGCGAGACCAGCTACGCGCAGAACTCCTCGCTTCAGAAGCGGGGCATGGACACGCTGAAGAGCCTTATCAC<br>CAACTCGGCGACGGACGTGTACATCGCGCAGATGCCGGAGAGGTTCACGGTGGCCGACCTGGGCTGCTCGTCGGGC<br>CCGAACGCGCTGTGCCTCGTCGAGGACATCGTCGGGAGCATCGGCCGGGTGTGCGGCCGGTCGTCGCAGCCGCCGC<br>CCGAGTTCTCGGTGCTCCTCAACGACCTCCCGACCAACGACTTCAATACCATCTTTTTCAGCCTGCCGGAGTTCAC<br>CGACCGGCTCAAGGCCGCCGCCGAGACCGATGAGTGGGGCCGGCCGATGGTGTTCCTGTCCGGCGTGCCGGGGTCC<br>TTCTACGGGAGGCTCTTCCCAAGGAAGAGCGTGCACTTCATCTGCTCCTACTCCAGCTTGCACTGGCTCTCCCAGG<br>TCCCGCCGGGGCTCTTCGACGAGGCCACGGGCACGCCCATCAACAAGGGGAAGATGTACATCTCGAGCTCCAGCCC<br>GCTCGCCGTGCCGACAGCCTACCTGAGGCAGTTCCAGAGGGACTTCGGCCTGTTCCTCAGATCACGCGCCGCCGAG<br>GTCGTCGCCGGCGGCCGGATGGTCCTCGCCATGCTCGGCAGGCAGACCGAGGGCTACATCGACAGGCGAACCACCT<br>TCCTCTGGGAGCTCCTCTCCGAGTCGTTCGCCTCGCTCGTGGCACAGGTGGGTGATCGAGCTTGCCATTCGAGAAG<br>GTGGACGCGTACAACGTGCCGTTTCTACGCGCCGTCGACCCAGCAGGTGGAGGAGGAGGTGCGGCGAGAGGGGTCG<br>TTCCGGCTCGACTACGTGCAGACGTACGAGATCAACCTGAGCAGCAGCGGTGACGCCAAGGAGGACGGCCGGACGG<br>TGTCCATGGCGATCAGGGCCATCCAGGAGTCCATGCTCAGCCACCACTTCGGCCCGGACATCGTCGATGCGCTCTT<br>CCACAGTAACGAGTACTGGTTCTGGATGTAA |

**Supplementary Table 7. Primers used in this study.**

| Purpose | Primer name | Sequence (5'-3') | Notes |
|---|---|---|---|
| Cloning | CLAMT1aF | ATGGCGTCCTCGCTGCTCCAC | |
| | CLAMT1aR | TCATAACCTTGTGAGGACGAC | |
| | CLAMT1a-bamH1F | ATTGGATCCATGGCGTCCTCGCTGCTCCAC | |
| | CLAMT1a-not1R | ATTGCGGCCGCTCATAACCTTGTGAGGACGAC | |
| | CLAMT1bF | ATGGCTTCTTCCTCCAATTGCTCC | |
| | CLAMT1bR | TCACAACAACCTTGTCAGGACCAC | |
| | CLAMT1b-Xba1F | AATTCTAGAATGGCTTCTTCCTCCAATTGCTCC | |
| | CLAMT1b-not1R | AATGCGGCCGCTCACAACAACCTTGTCAGGACCAC | |
| | CLAMT1cF | ATGGCGTCCTCGCTGCTCCAC | |
| | CLAMT1ciso1R | CACAGGAAACAGCTATGACC | |
| | CLAMT1ciso2R | TTACATCCAGAACCAGTACTCG | |
| | CLAMT1c-bamH1F | ATTGGATCCATGGCGTCCTCGCTGCTC | |
| | CLAMT1ciso1-Not1R | AATGCGGCCGCCACAGGAAACAGCTATGACC | |
| | CLAMT1ciso2-Not1R | AATGCGGCCGCTTACATCCAGAACCAGTACTCG | |
| | AtCLAMTF | ATGGATAAGAAGGATATGGAG | |
| | AtCLAMTR | TCAGAGCTTCTTTCTTAGGAC | |
| | AtCLAMT-BamH1F | ATTGGATCCATGGATAAGAAGGATATGGAG | |
| | AtCLAMT-Not1R | ATTGCGGCCGCTCAGAGCTTCTTTCTTAGGAC | |
| | AtMAX1-BamH1F | AATGGATCCATGAAGACGCAACATCAATGGT | |
| | AtMAX1-Not1R | AATGCGGCCGCTCAGAATCTTTTGATGGTTCTGAGCT | |
| Genotyping | CLAMT1bF | CACAGCCAAGCTGAAAGCCG | |
| | CLAMT1bR | CGTGCTTGATATGAACATCTTCCC | |
| | CLAMT1cF | CGACTTCAATACCATCTTTTTCAGC | |
| | CLAMT1cR | CCAGTGCAAGCTGGAGTAGG | |
| | UBC-E2F | ACCGCCTGACAATCCCTATG | Shivhare and Lata, 2016 |
| | UBC-E2R | GGGATAGTCTGGCGGAAAATG | Shivhare and Lata, 2016 |
| RT-PCR | *EcPT4 F* | ACGCCTACGACCTATTCTGCATCACC | Pudake et al., 2017 |
| | *EcPT4 R* | GCCGAACACGAG CTGGCCTATCA | Pudake et al., 2017 |
| | *RiEFαf* | GCTATTTTGATCATTGCCGCC | Fiorilli et al., 2016 |
| | *RiEFαr* | TCATTAAAACGTTCTTCCGACC | Fiorilli et al., 2016 |
| | *RiPEIP1* | AAGAAAGTAAACGTGTGGCT | Fiorilli et al., 2016 |
| | *RiPEIP1* | TAACACTCATCTCGGGACTG | Fiorilli et al., 2016 |
| | *SiPHT1;9 F* | CAAGGAGATAAACGCCCTGAC | Ceasar et al., 2014 |
| | *SiPHT1;9 R* | ACCGATGAGCTGGATCAGGTA | Ceasar et al., 2014 |
| | *TUA F* | GAGCGTCTGTCTGTTGACTATG | Saha et al., 2014 |
| | *TUA R* | GTGGACAGGACACTGTTGTATG | Saha et al., 2014 |
